# The transcription factor Hhex cooperates with the corepressor Tle3 to promote memory B cell development

**DOI:** 10.1101/2020.02.25.965301

**Authors:** Brian J. Laidlaw, Lihui Duan, Ying Xu, Sara E. Vazquez, Jason G. Cyster

## Abstract

Memory B cells (MBCs) are essential for long-lived humoral immunity. However, the transcription factors (TFs) involved in MBC differentiation are poorly defined. Here, by single cell RNAseq analysis, we identified a population of germinal center (GC) B cells in the process of differentiating into MBCs. Using an inducible Crispr/Cas9 screening approach we identified the hematopoietically-expressed homeobox gene Hhex as a transcription factor regulating MBC differentiation. The co-repressor Tle3 was also identified in the screen and was found to interact with Hhex. Bcl6 directly repressed *Hhex* in GC B cells. Reciprocally, Hhex-deficient MBCs exhibited derepressed *Bcl6* and reduced expression of Bcl6-repressed *Bcl2*. Overexpression of Bcl2 was able to rescue MBC differentiation in Hhex-deficient cells. We also identified Ski as an Hhex-induced transcription factor involved in MBC differentiation. These findings establish an important role for Hhex-Tle3 in regulating the transcriptional circuitry governing MBC differentiation.

## INTRODUCTION

Memory B cells (MBCs) are long-lived cells that mediate humoral immunity by undergoing rapid proliferation and differentiation into antibody-secreting cells following antigen reencounter. MBCs are superior to long-lived plasma cells in responding to variant strains of pathogens such as West Nile virus and influenza ^1, 2^. While MBCs can arise independently of the germinal center (GC), the GC is necessary for the affinity maturation of MBCs ^3^. The isotype class-switched MBC response to T-dependent antigens is largely GC-derived ^4, 5^.

Understanding how GC B cells differentiate into MBCs is important for the development of therapeutics capable of modulating MBC development. Multiple models have been postulated to explain MBC development. The stochastic model suggests that MBCs are randomly selected from GC B cells and is supported by data from various genetic models showing that a decline in GC B cells often results in a proportional loss in MBCs ^6–8^. The instructive model posits that MBC development is actively regulated by cell extrinsic signals such as cytokines and cell-contact dependent signals. This model is supported by the finding that perturbations in the ability of B cells to receive T cell help through ablation of the ability to sense IL-21 or migrate to the dark zone of the GC results in an accumulation of MBCs disproportionate to GC size ^9–11^. Additionally, follicular helper T (T_FH_) cell-derived IL-9 can promote MBC development in some settings ^12, 13^. MBCs tend to emerge from the GC prior to plasma cells and are predominantly derived from lower affinity GC B cells that receive less T cell help ^14, 15^. Low T cell help is associated with elevated expression of Bach2, which is a transcription factor (TF) that predisposes GC B cells to develop into MBCs ^14^. The majority of GC B cells express Bach2, indicating that other transcriptional regulators are necessary to guide MBC development ^14^.

We have previously characterized a population of GC B cells differentiating into MBCs following viral infection ^16^. GC memory precursor (PreMem) B cells are transcriptionally and functionally distinct from GC cells and are localized near the GC border ^12, 16^. PreMem B cells are found in both mice and humans ^17^. In this study we sought to identify transcriptional regulators of MBC development. Using a CRISPR/Cas9-based knockdown screen we found that ablation of the hematopoietically expressed homeobox protein Hhex or the corepressor Tle3 (transducin-like enhancer of split-3) resulted in impaired MBC differentiation. Hhex has been reported to interact with Tle1 ^18^ and we find that Hhex directly interacts with Tle3. We determined that Hhex is necessary for the development of PreMem B cells and that Bcl6 represses *Hhex* expression in GC B cells. Hhex-deficient MBCs have reduced expression of the Bcl6 target gene *Bcl2*, and overexpression of Bcl2 is sufficient to rescue MBC differentiation in Hhex-deficient cells. The Hhex-induced TF Ski also promoted MBC differentiation. Together our data indicates that Hhex cooperates with Tle3 to regulate the transcriptional circuitry governing GC B cell differentiation into MBCs.

## RESULTS

### Identification of PreMem B cells using single cell RNA sequencing

We performed droplet-based single cell RNA sequencing (scRNA-seq) on sorted splenic B cells from mice at day 11 post acute lymphocytic choriomeningitis virus strain Armstrong (LCMV) infection. The sorted population consisted of a mixture of 90% GC B cells, 5% follicular (FO) B cells, and 5% MBCs (**gating shown in Fig. S1a**). Day 11 post infection (p.i.) corresponds to the peak of PreMem B cells following LCMV infection ^16^. After quality control, we retained 11,238 cells with an average of 1,574 genes per cell; 98.9% of cells were retained for further analysis after removal of non-B cells.

Unsupervised clustering using Seurat ^19^ revealed 8 clusters, as visualized using tSNE, with clusters ranging in size from 207 to 2,765 cells with 33-417 differentially expressed genes (DEGs) per cluster (**Fig. 1a, b**). The specific DEGs expressed by each cluster (**Fig. 1c, Fig. S1b, c**) suggested assignment to specific cell states, namely FO, GC LZ, GC DZ and MBC, based on previously described gene expression profiles (**Fig.1a**). Validation of assignment to specific cell states was performed by assigning cells with a score corresponding to their similarity with gene expression profiles distinguishing these states based on bulk RNAseq analysis (**Fig. S1d**) ^16, 20^. Cells were also assigned to a specific cell cycle stage based on their gene expression profile (**Fig. S1d**). The identification of a Myc^+^ GC cluster is consistent with work identifying ∼5% of GC B cells as expressing Myc ^21^. The significance of the subclustering of LZ and DZ states was not a focus of the current studies and merits future investigation.

**Figure 1.**
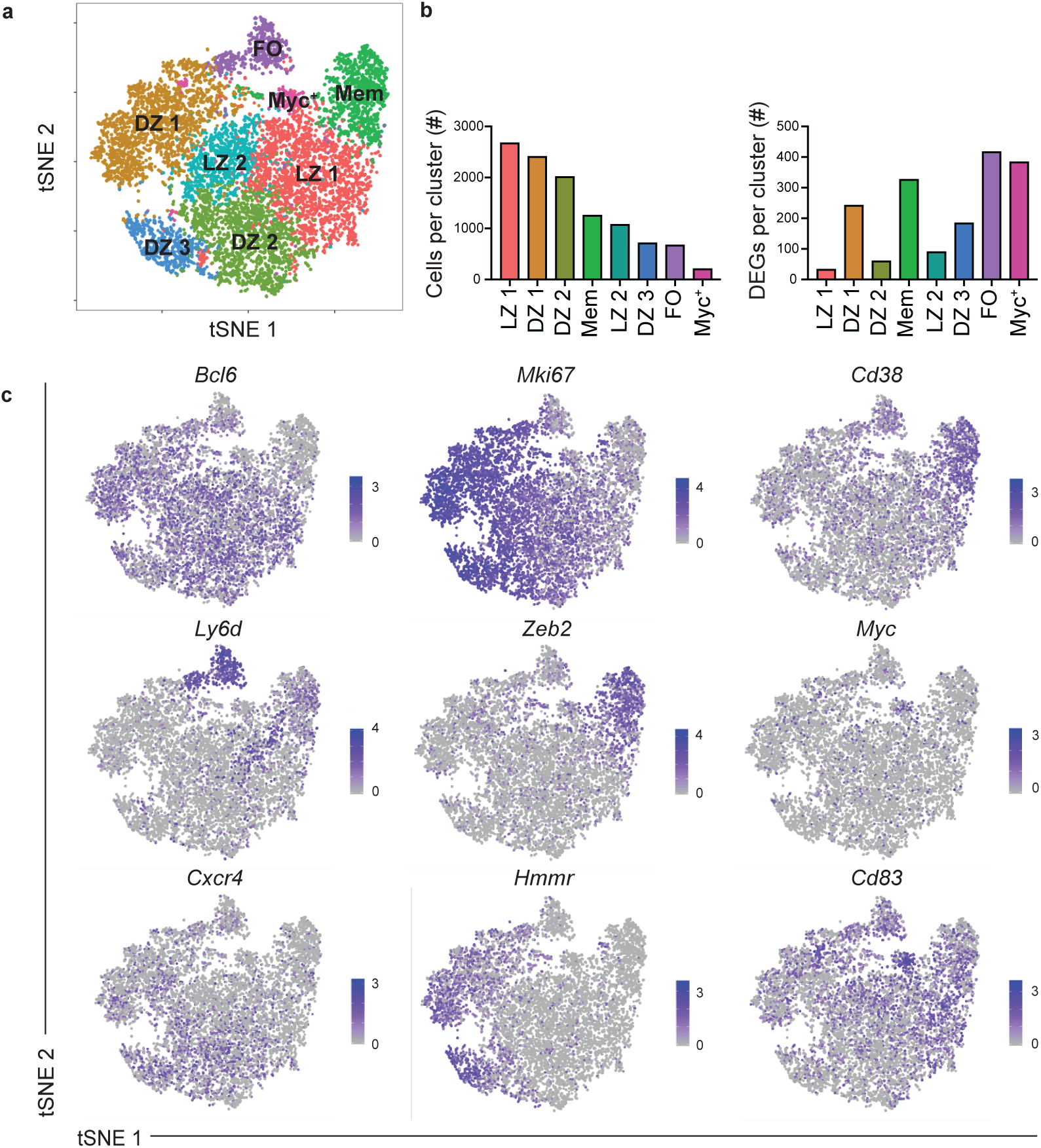
Identification of eight B cell clusters following viral infection using scRNAseq. (**a**) Unsupervised clustering of splenic B cells at day 11 post LCMV infection visualized with tSNE. Each point is a single cell colored by cluster assessment. The sorted population consisted of a mixture of 90% GC B cells, 5% FO B cells, and 5% MBCs (**b**) Number of cells per cluster (left) and DEGs per cluster (right). (**c**) Gene expression distinguishing the eight clusters projected onto tSNE plots. Color scaled for each gene with highest log-normalized expression level noted. See also Figure S1.

Among the identified clusters was a population of cells that transcriptionally resembled MBCs, labeled as Mem (**Fig. 1a**). As PreMem B cells have a similar transcriptional profile to MBCs, we performed additional subclustering analysis of the Mem cluster in order to distinguish the PreMem B cell population ^16^. We found that the Mem cluster could be divided into 3 clusters, including a population of PreMem B cells and 2 populations of MBCs (**Fig. 2a**). Cluster identity was determined based on similarity to gene expression profiles distinguishing MBCs, GC B cells, and PreMem B cells (**Fig. 2b**). The PreMem cluster consisted of 106 cells with 257 DEGs relative to other cells present in the Mem cluster (**Fig. S2a**). PreMem B cells expressed genes associated with both the GC state (*Mki67*, *Bcl6*, *Fas*) and the MBC state (*Cd38*, *Sell*, *Bcl2*) supporting the notion that PreMem cells represent a transitional population (**Fig. 2c, d**). While both the Mem 1 and Mem 2 clusters expressed MBC-associated genes, Mem 1 cells displayed higher expression of many MBC-associated genes and had the lowest expression of *Mki67* (**Fig. 2d**). Pseudotime trajectory analysis using Monocle ^22^ showed that Mem 1 cells are enriched in the branch furthest from where PreMem cells are most abundant, further indicating that the Mem 1 cluster represents more quiescent MBCs (**Fig. S2b**). The Mem 2 cluster is enriched in branches closer in proximity to PreMem cells suggesting that Mem 2 cells have more recently emerged from the GC (**Fig. S2b**).

**Figure 2.**
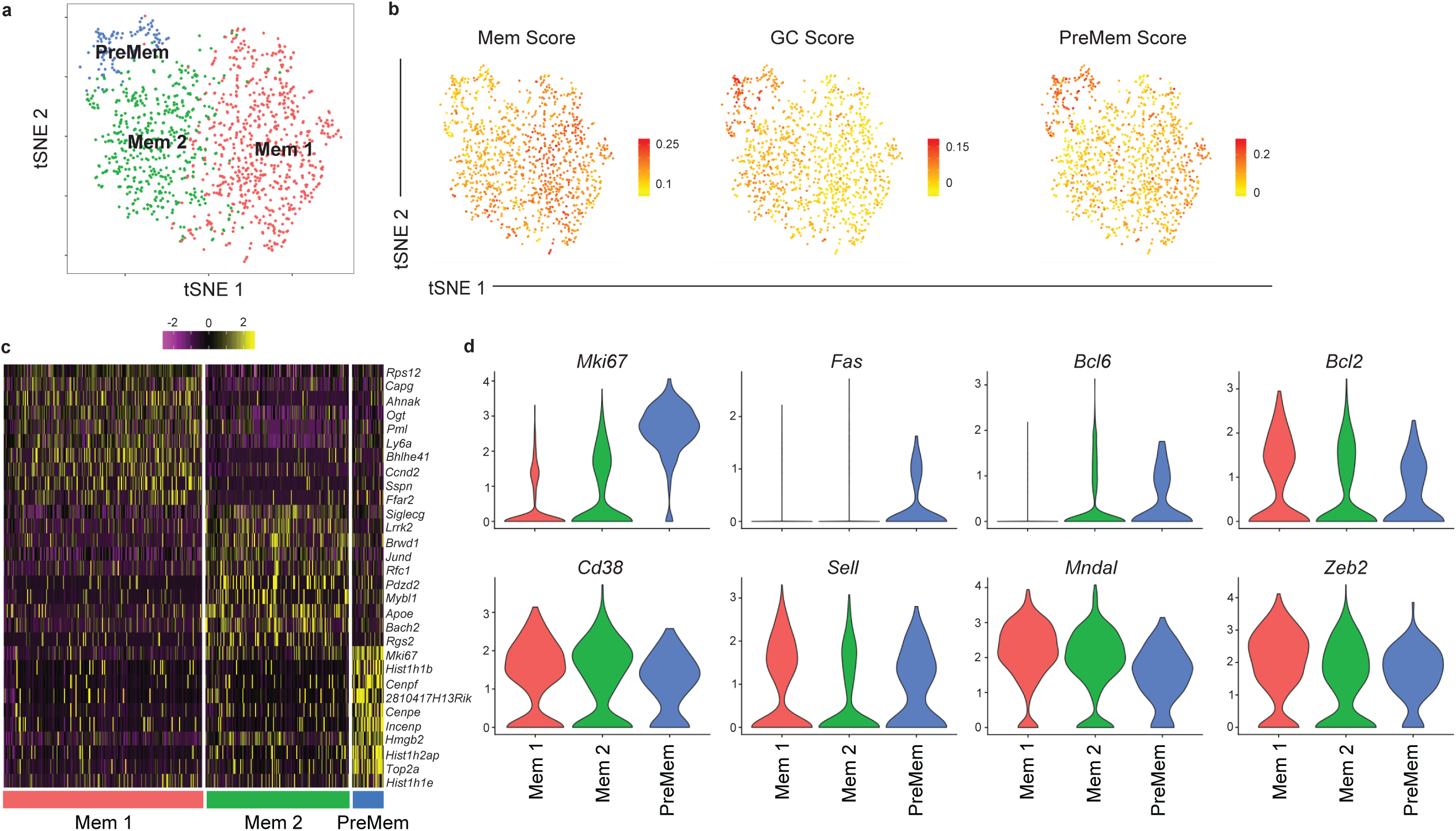
Identification of PreMem B cells using scRNAseq. (**a**) Subclustering analysis of the Mem cluster of splenic B cells at day 11 post LCMV infection visualized with tSNE. Each point is a single cell colored by cluster assessment. (**b**) Enrichment score for gene signatures distinguishing MBCs (left), GC B cells (middle), and PreMem B cells (right) projected onto tSNE plots. Color scaled for each gene with highest log-normalized expression level noted. (**c**) Heatmap of each cell’s (column) expression of the top ten DEGs per cluster (rows). Log-normalized expression scaled for each gene. Cluster name displayed below. (**d**) Violin plots of select gene expression by cluster with highest log-normalized expression value labeled. See also Figure S2.

### Conditional Crispr-Cas9 screen of transcription factors expressed by PreMem B cells

MBCs displayed elevated expression of a number of TFs relative to GC B cells (**Fig. 3a**). Importantly, many of these TFs are already expressed in PreMem B cells based on single cell and bulk RNAseq analysis (**Fig. S3a, b**). Human MBCs also display elevated expression of the same TFs relative to GC B cells (**Fig. S3c**) ^23^. We sought to perform a screen of TFs regulating MBC development. As FO B cells often express TFs expressed by MBCs, we used a conditional Cas9 approach to specifically ablate TF expression in activated B cells. We intercrossed Rosa26-LSL-Cas9 and *Cg1*^Cre^ mice to generate mice in which Cas9 activity is limited to cells expressing endogenous *Ighg1*, thereby strongly enriching for Cre activity among GC-derived cells ^24, 25^.

**Figure 3.**
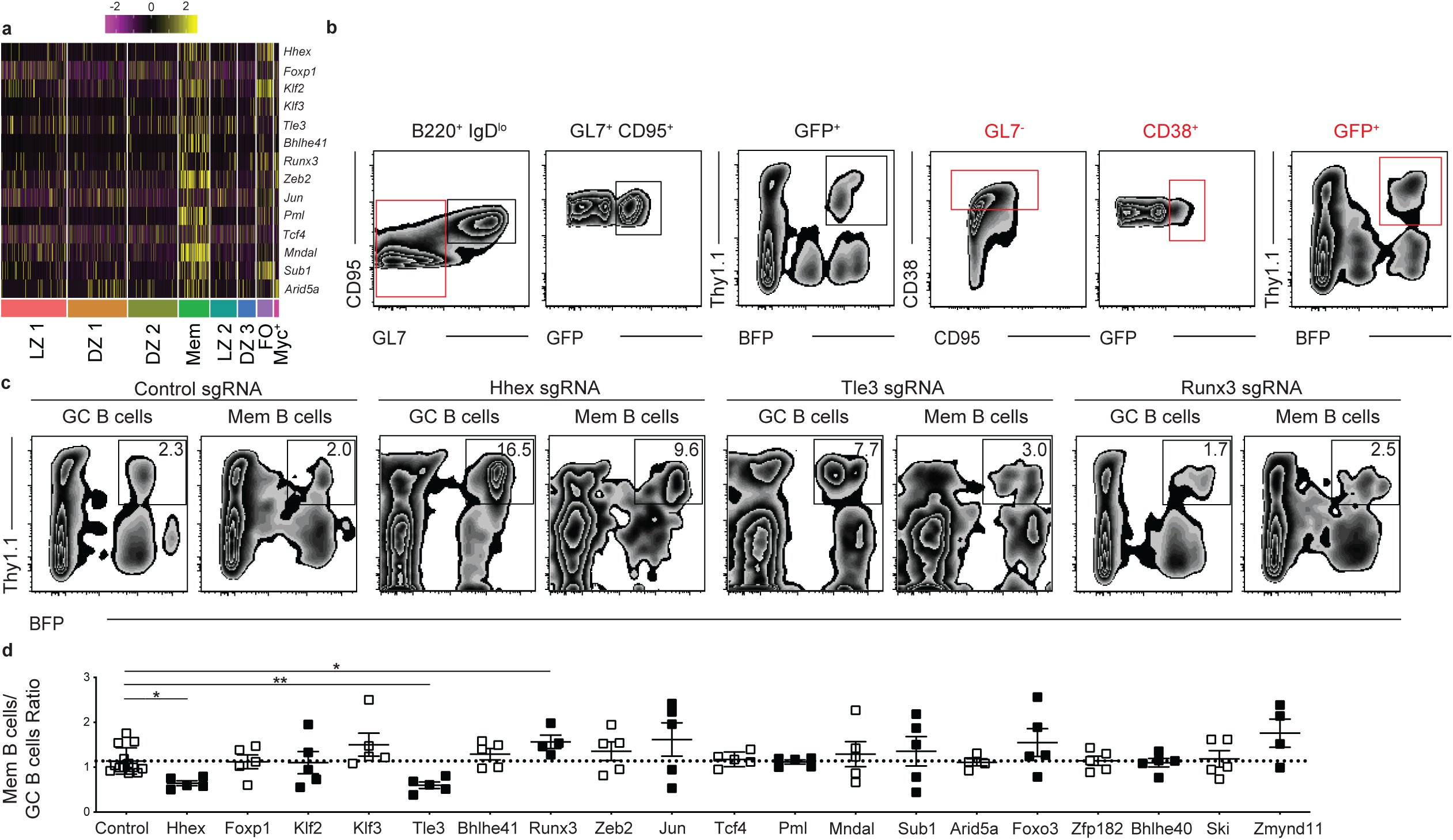
*In vivo* conditional Cas9 screen of transcription factors expressed by PreMem B cells. (**a**) Heatmap of each cell’s (column) expression of select TFs per cluster (rows). Log-normalized expression scaled for each gene. Cluster name displayed below. (**b**) Gating strategy for conditional Cas9 screen. Mice were sacrificed at day 30 p.i. and splenic B cells were analyzed as shown. Gates are color coded and shown in series from left to right. (**c**) Representative FACS plots showing the percentage of Thy1.1^+^BFP^+^ GC and MBCs at day 30 post LCMV infection in mice containing cells transduced with sgRNAs targeting a control nonexpressed gene, *Hhex*, *Tle3*, or *Runx3*. Transduction efficiencies varied between constructs and sets of BM chimeras. (**d**) Ratio of Thy1.1^+^BFP^+^ MBCs to Thy1.1^+^BFP^+^ GC B cells for mice containing cells transduced with sgRNA targeting the listed genes. Data are pooled from 5 independent experiments with 3-5 mice per group. Statistical analyses were performed using the ordinary one-way ANOVA with Dunnett multiple comparison testing (*, *p* < 0.05; **, *p* < 0.01). See also Figure S3.

Bone marrow from Rosa26-LSL-Cas9^f/+^ *Cg1*^Cre/+^ mice was transduced with two single guide (sg) RNA-containing retroviral constructs per gene target (expressing either Thy1.1 or BFP as reporters) and then transferring to lethally irradiated recipient mice. Following reconstitution, mice were infected with LCMV and analyzed on day 30 p.i. The percentage of Thy1.1^+^BFP^+^ cells among the Cre expressing GC and MBC population was then determined (**Fig. 3b**). The efficiency of this approach was tested using sgRNAs specific for CD38 and CXCR3 with it being found that there was 70-99% ablation of target protein expression on Cre-expressing B cells as quantified by flow cytometric analysis (**Fig. S3d, e**). We screened 19 TFs expressed by MBCs and identified 3 TFs for which gene ablation led to a significant change in the fraction of MBCs relative to GC B cells compared to control sgRNA transduced cells (**Fig. 3c, d**). Specifically, we found that ablation of Hhex or Tle3 led to impaired MBC development, while loss of Runx3 resulted in a slight increase in MBC development (**Fig. 3c, d**).

### Retroviral overexpression of Hhex or Tle3 promotes memory B cell development

We next investigated whether retroviral overexpression of Hhex or Tle3 was sufficient to promote MBC development. Bone marrow was transduced with plasmids expressing these TFs and then transferred to lethally irradiated recipient mice. Following reconstitution, mice were infected with LCMV and analyzed on day 30 p.i. The percentage of transduced FO, GC, and MBCs was then determined. We found that overexpression of Hhex or Tle3 was sufficient to promote MBC development relative to the FO or GC B cells (**Fig. 4a, b**). Overexpression of Hhex or Tle3 also led to a decrease in the percentage of GC B cells relative to FO B cells (**Fig. 4a, b**). The increased fraction of Hhex-overexpressing MBCs relative to GC B cells was consistent between days 15-60 p.i. (**Fig. S4a**). Hhex overexpression also promoted the development of NP^+^ MBCs following immunization with NP-CGG in alum, indicating that Hhex can promote MBCs during both Th1-type and Th2-type immune responses (**Fig. S4b, c**).

**Figure 4.**
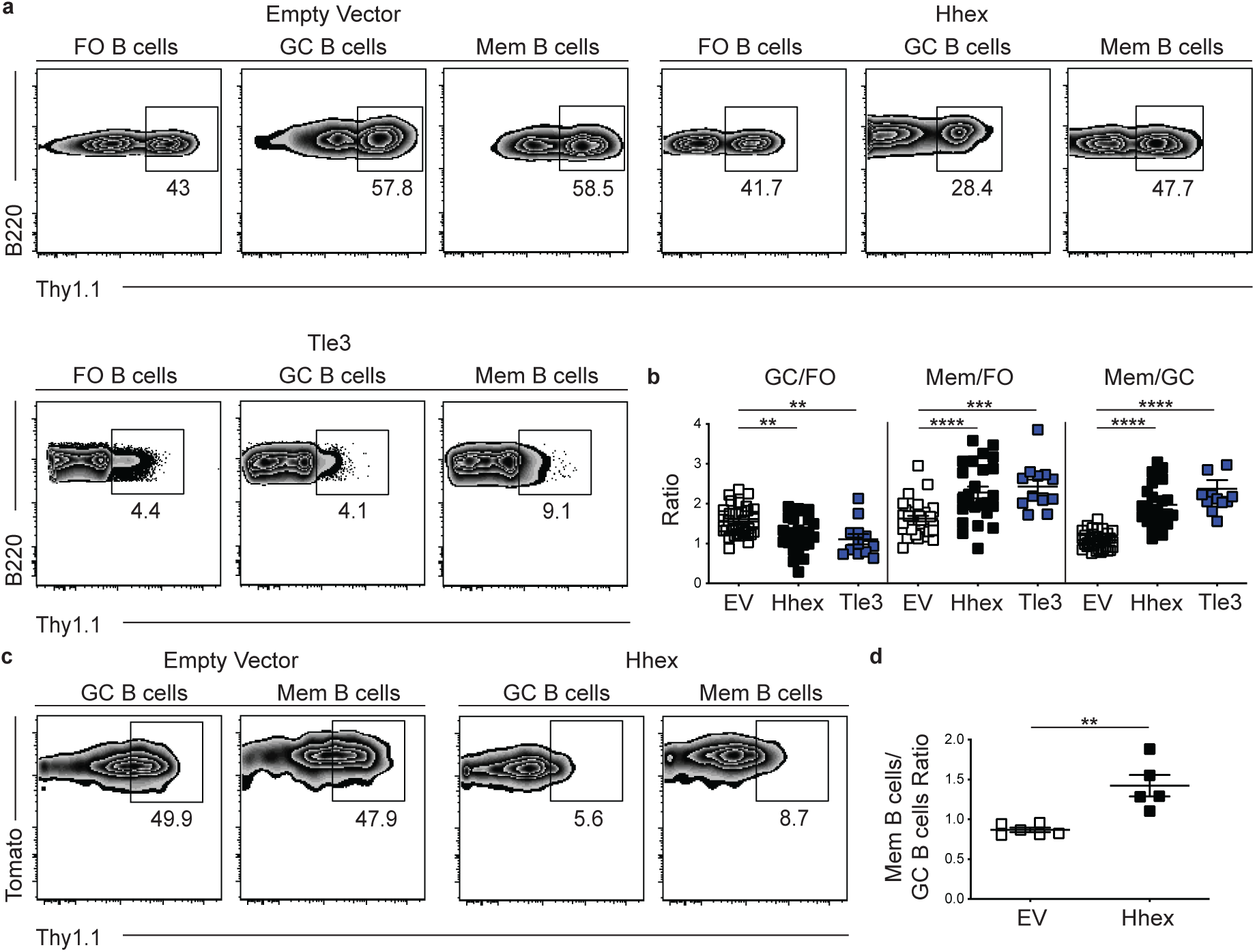
Overexpression of Hhex or Tle3 promotes memory B cell differentiation. (**a**) Representative FACS plots of the percentage of transduced (Thy1.1^+^) cells among splenic FO (B220^+^IgD^hi^GL7^-^CD38^+^CD95^-^), GC (B220^+^IgD^lo^GL7^+^CD95^+^), and MBCs (B220^+^IgD^lo^GL7^-^ CD38^+^CD95^+^CD73^+^) at day 30 post LCMV infection with empty vector, Hhex, or Tle3. Transduction efficiencies varied between constructs. (**b**) Ratio of transduced GC to FO B cells (left), MBCs to FO B cells (middle), and MBCs to GC B cells (right) in experiments of the type in a. Data are pooled from 7 independent experiments with 3-6 mice per group. (**c**) Representative FACS plots of the percentage of transduced (Thy1.1^+^) cells among splenic GC (B220^+^IgD^lo^GL7^+^CD95^+^Tomato^+^), and MBCs (B220^+^IgD^lo^GL7^-^CD38^+^Tomato^+^) at day 30 post LCMV infection of mice reconstituted with S1pr2-ERT2cre TdTomato bone marrow transduced with a control (EV) or Hhex expressing loxp-EGFP-loxp-MSCV-IRES-Thy1.1 vector. Mice were treated with Tm at day 4 p.i. to induce Cre expression. (**d**) Ratio of transduced MBCs to GC B cells in the experiment described in c. Data are representative of 2 independent experiments with 5-6 mice per group. Statistical analyses were performed using the unpaired two-tailed Student’s *t*-test or the ordinary one-way ANOVA with Dunnett multiple comparison testing (*, *p* < 0.05; **, *p* < 0.01; ***, *p* < 0.001; ****, *p* < 0.0001). See also Figure S4.

We tested several additional TFs using the retroviral overexpression approach and found that Mndal, Zmynd11, and Klf3 overexpression failed to significantly promote MBC development (**Fig. S5a, b**). Interestingly, overexpression of Ski or Klf2 was sufficient to promote MBC development relative to GC B cells (**Fig. S5a, b**). These findings may be in accord with the observation that retroviral overexpression of Ski and Klf2 imparts a competitive advantage to CD40-engaged B cells *in vitro* ^26^.

To further explore the role of Hhex in MBCs we tested whether retroviral overexpression specifically in GC B cells was sufficient to promote MBC development. To perform this experiment, we generated a loxp-EGFP-STOP-loxp-MSCV-IRES-Thy1.1 vector in which *Hhex* was inserted downstream of the floxed EGFP-STOP element. Expression of Cre recombinase by cells transduced with this vector will lead to excision of the loxp-EGFP-STOP-loxp sequence and subsequent expression of the downstream gene. Bone marrow from S1pr2-ERT2creTdTomato mice was transduced with an *Hhex* expressing loxp-EGFP-STOP-loxp-MSCV-IRES-Thy1.1 plasmid and transferred to lethally irradiated recipient mice ^14^. *S1pr2* is highly and specifically expressed in GC B cells so Cre expression among B cells in S1pr2-ERT2creTdTomato mice is restricted to GC-derived cells ^27^. Following reconstitution, mice were infected with LCMV, treated with tamoxifen (Tm) beginning at day 4 to induce Cre expression in *S1pr2* expressing cells, and analyzed on day 30 p.i. We found that Hhex overexpression promoted MBC development relative to empty vector transduced cells (**Fig. 4c, d**). This indicates that Hhex can specifically function in GC B cells to promote their differentiation into MBCs.

### Hhex promotes MBC development through direct binding to DNA

Hhex is a homeodomain TF that is important for the development of common lymphoid progenitor cells and which promotes stem cell self-renewal under conditions of hematopoietic stress ^28, 29^. Hhex consists of three functional domains capable of contributing to the regulation of gene expression (**Fig. 5a**) ^30^. To determine which domains contribute to the promotion of MBC development, we generated retroviral constructs containing mutant forms of Hhex that either lack the ability to bind DNA (N188A) or the acidic C-terminal activation region (ΔCT), which is involved in transcriptional activation of Hhex-regulated genes ^31, 32^. We found that cells overexpressing the Hhex N188A mutant had reduced MBC development relative to EV-transduced FO or GC B cells (**Fig. 5b, c**). Cells overexpressing HhexΔCT had similar MBC development to EV-transduced cells, but did not display the enhanced MBC development evident in cells overexpressing full length Hhex overexpression (**Fig. 5b, c**). These data indicate that the ability of Hhex to promote MBC development is dependent on both binding DNA via the homeodomain and on the C-terminal domain.

**Figure 5.**
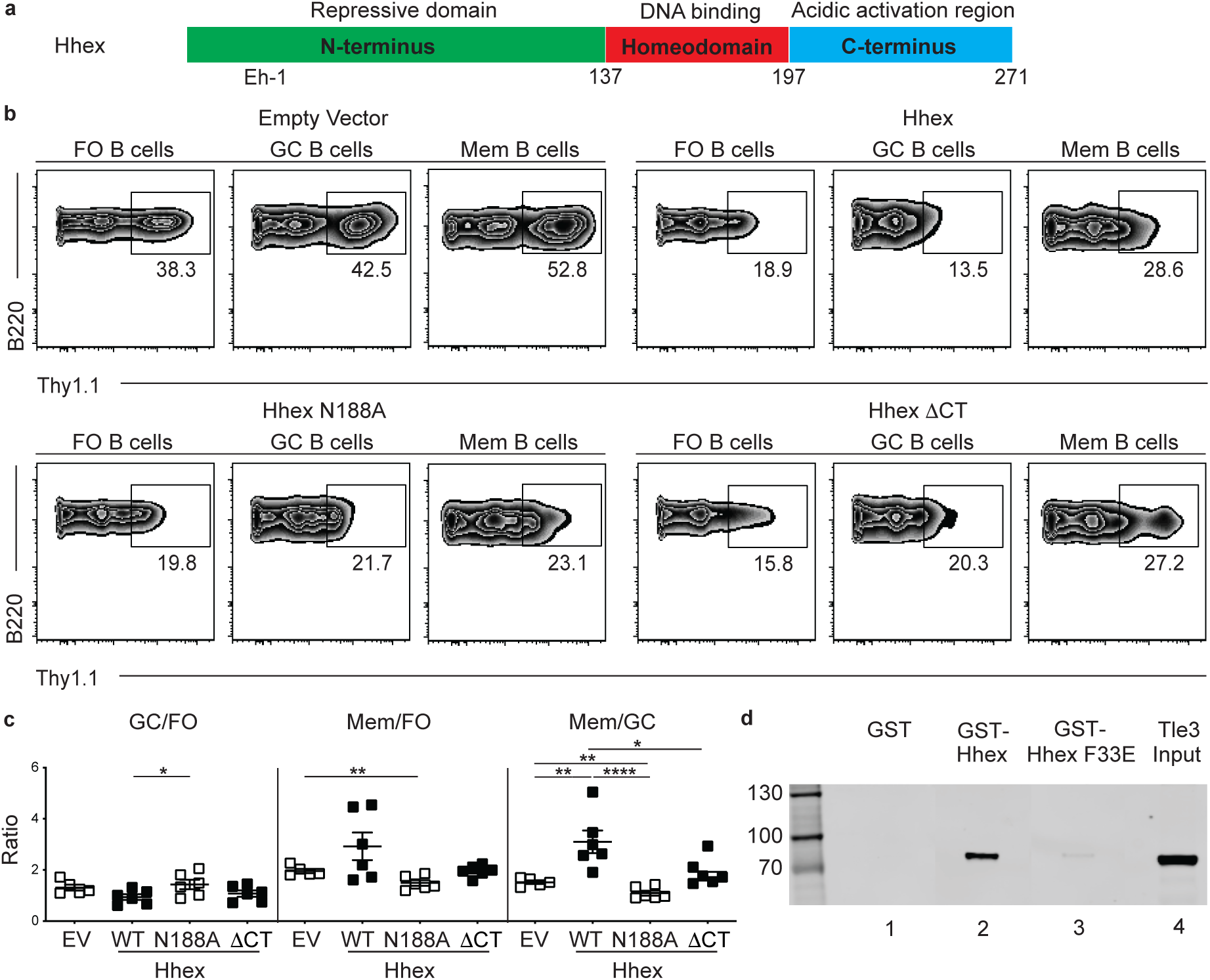
Hhex interacts with Tle3 and promotes memory B cell differentiation through binding to DNA. (**a**) Schematic representation of the domains found in Hhex. Location of Eh-1 motif (aa 31-41, TPFYIDDILGR) in N-terminus is indicated. (**b**) Representative FACS plots of the percentage of transduced cells among splenic FO (B220^+^IgD^hi^GL7^-^CD38^+^CD95^-^), GC (B220^+^IgD^lo^GL7^+^CD95^+^), and MBCs (B220^+^IgD^lo^GL7^-^CD38^+^CD95^+^CD73^+^) at day 30 post LCMV infection in empty vector, Hhex, Hhex N188A, and Hhex ΔCT-overexpressing bone marrow chimeras. (**c**) Ratio of transduced GC to FO B cells (left), MBCs to FO B cells (middle), and MBCs to GC B cells (right). Data are representative of 2 independent experiments with 4-6 mice per group. Statistical analyses were performed using the unpaired two-tailed Student’s *t*-test or the ordinary one-way ANOVA with Dunnett multiple comparison testing (*, *p* < 0.05; **, *p* < 0.01; ****, *p* < 0.0001). (**d**) Interaction of in vitro translated and transcribed Tle3-FLAG with glutathione beads coated with GST (lane 1), GST-Hhex (lane 2), or GST-Hhex F33E (lane 3) detected by FLAG Western blot. Lane 4 is a control showing 20% of Tle3 input. Tle3 is a 772aa protein with an MW of 83kDa. See also Figure S5.

### Hhex directly interacts with Tle3

Tle3 is a member of the Tle/Groucho family of co-repressors. These proteins do not have a DNA binding domain but interact with multiple transcription factors through either of two conserved domains ^33^. Hhex was found to interact via a conserved N-terminal Eh-1 motif with Tle1 ^18^. This motif was originally identified in *Drosophila* homeodomain proteins Engrailed and Goosecoid, where it mediates their interaction with Groucho ^34^. Mutation of the Eh-1 motif in human Hhex (F32E) disrupted Tle1 binding and prevented Hhex-mediated gene repression ^18^. Given that both Hhex and Tle3 promoted MBC development, we sought to determine whether Hhex interacted with Tle3. We generated gluthathione S-transferase (GST) fusion proteins expressing either the wild type N-terminal domain of mouse Hhex or a version with a mutation in the Eh-1 motif (F33E) (**Fig. S5c**) and used them in binding assays with *in vitro* translated C-terminal Flag-tagged Tle3. Tle3 was found to bind strongly to GST-Hhex but weakly, if at all, to GST-Hhex F33E or GST alone (**Fig. 5d**). Similar findings were made in pull down assays with radiolabeled Tle3 (**Fig. S5d**). Neither GST-Hhex or GST-Hhex F33E displayed binding to TFs not known to interact with Hhex (**Fig. S5e**). These data suggest that Hhex and Tle3 cooperate to promote MBC development.

### Inducible ablation of Hhex leads to reduced MBC development

To further explore how Hhex ablation affects MBC development, we intercrossed *Hhex*^flox/flox^ and S1pr2-ERT2creTdTomato mice. We then infected *Hhex*^+/+^ (WT), *Hhex*^f/+^ (Het), or *Hhex*^f/f^ (KO) S1pr2-ERT2creTdTomato mice with LCMV and treated with Tm beginning at day 4 p.i (**Fig. 6a**). Mice were analyzed at day 30 p.i. and we found that cells lacking Hhex displayed a marked reduction in the percentage and number of MBCs (**Fig. 6b-d**). There was no difference in the number or percentage of GC B cells in Hhex-deficient mice (**Fig. 6c-d**). A reduction in the magnitude of the NP-specific MBC response was also found following NP-CGG in alum immunization of Hhex KO S1pr2-ERT2creTdTomato mice (**Fig. S6a-c**). The relative reduction in the percentage of MBCs in Hhex-deficient mice was similar at days 15 and 45 post LCMV infection, with no difference in the percentage of GC B cells apparent at either time point (**Fig. 6e**). Loss of Hhex did not impair the development of GC-derived CD138^+^ plasma cells or LCMV-specific IgG antibodies (**Fig. 6f, g**). We also did not observe differences in the proportion of proliferating or apoptotic GC cells in Hhex-deficient mice, as indicated by the percentage of BrdU^+^ or Casp3^+^ GC B cells respectively (**Fig. 6h**).

**Figure 6.**
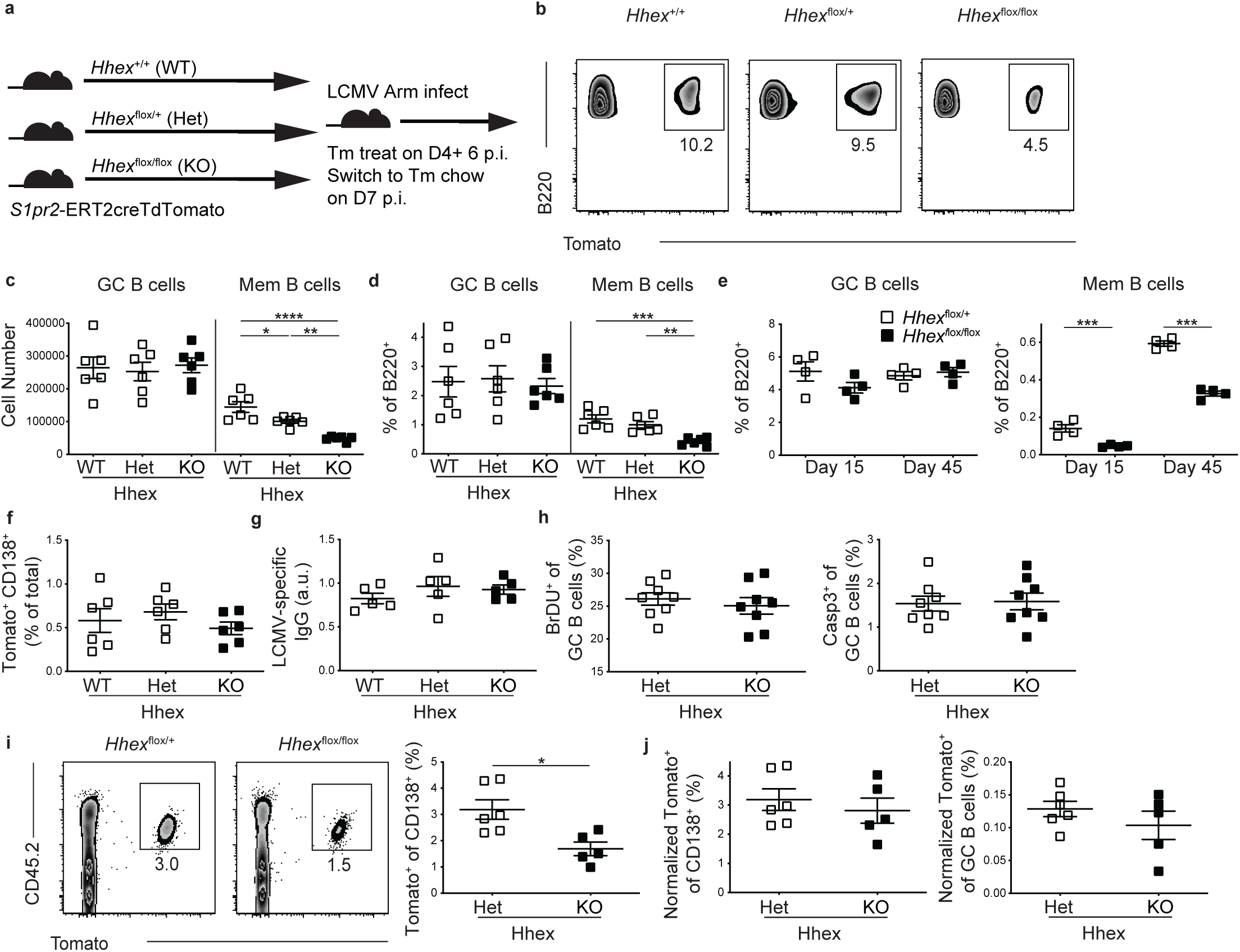
Ablation of Hhex in GC B cells impairs memory B cell differentiation. (**a**) Experimental schematic. *Hhex*^+/+^ (WT), *Hhex*^flox/+^ (Het), and *Hhex*^flox/flox^ (KO) S1pr2-ERT2cre TdTomato mice were infected with LCMV, treated with Tm beginning at day 4, and then analyzed at day 30 p.i. (**b**) Representative FACS plots of the percentage of Tomato^+^ cells among splenic B220^+^IgD^lo^GL7^-^CD38^+^ MBCs in the groups indicated in a. (**c, d**) Number (c) or percentage (d) of GC B cells (B220^+^IgD^lo^GL7^+^CD95^+^Tomato^+^) and MBCs (B220^+^IgD^lo^GL7^-^ CD38^+^Tomato^+^) in Hhex WT, Het, and KO mice at day 30 post LCMV infection. Data in a-d are representative of 4 independent experiments with 4-6 mice per group. (**e**) Percentage of B cells that are GC B cells (left) and MBCs (right) in Hhex Het and KO mice at days 15 and 45 post LCMV infection. Data are representative of 2 independent experiments with 4-6 mice per group. (**f**) Percentage of live splenic cells that are CD138^+^Tomato^+^ in Hhex WT, Het, and KO mice at day 30 post LCMV infection. (**g**) Quantification of anti-LCMV IgG in the serum of Hhex WT, Het, and KO mice at day 30 post LCMV infection. Data in f-g are representative of 2 independent experiments with 4-6 mice per group. (**h**) Percentage of GC B cells that were BrdU^+^ (left) and Casp3^+^ (right) in Hhex Het and KO mice at day 30 post LCMV infection. Data are pooled from 2 independent experiments with 4 mice per group. (**i**) Representative FACS plot (left) and percentage (right) of CD138^+^ cells that are CD45.2^+^Tomato^+^ at day 5 post challenge with LCMV. Equivalent numbers of splenic CD45.2^+^ T and B cells from Hhex Het and KO LCMV immune mice at day 30 p.i. were transferred to naïve CD45.1^+^ recipients one day prior to challenge. (**j**) Percentage of CD138^+^ cells (left) and GC B cells (right) that were CD45.2^+^Tomato^+^ at day 5 post challenge with LCMV when normalized to the percentage of MBCs present in the transferred cells. Data are pooled from 2 independent experiments with 3 mice per group. Statistical analyses were performed using the unpaired two-tailed Student’s *t*-test or the ordinary one-way ANOVA with Dunnett multiple comparison testing (*, *p* < 0.05; **, *p* < 0.01; ***, *p* < 0.001; ****, *p* < 0.0001). See also Figure S6.

To examine the functionality of Hhex-deficient MBCs, we transferred an equal number of B and T cells from Hhex Het and KO LCMV immune mice into congenically mismatched naive recipients and challenged the next day with LCMV. B cells transferred from Hhex KO mice displayed an impaired ability to differentiate into CD138^+^ cells following viral challenge (**Fig. 6i**). The magnitude of impairment corresponded to the reduced frequency of MBCs in the Hhex-deficient cells, as there was no difference in the ability of transferred cells to differentiate into CD138^+^ cells or GC B cells when normalized to the starting percentage of MBCs (**Fig. 6j**). Similar results were found using an analogous approach to assess MBC recall following NP-CGG in alum immunization (**Fig. S6d, e**). Together these data indicate that while Hhex is necessary for the development of MBCs during both Th1-type and Th2-type immune responses, it is not required for MBCs to respond upon antigen reencounter.

### Bcl6 represses expression of Hhex in GC B cells

We next examined how *Hhex* expression is regulated in GC B cells. *Hhex* is highly expressed in both FO and MBCs, but is downregulated in GC B cells (**Fig. 3a**). Bcl6 is highly expressed in GC B cells and serves to maintain GC B cell identity through repression of target genes. Analysis of publicly available chromatin immunoprecipitation (ChIP) sequencing data of human GC B cells ^35^ revealed that Bcl6 binds to the *Hhex* promoter (**Fig. 7a**). This binding occurs in complex with the corepressor BCOR, but not the corepressor SMRT (**Fig. 7a**). Retroviral overexpression of Bcl6 was sufficient to repress *Hhex* expression in mouse B cells (**Fig. 7b**). Reciprocally, retroviral overexpression of Hhex led to a reduction in GC B cells (**Fig. 4a, b**). Together, these data indicate that Bcl6 and Hhex function in opposition to regulate GC B cell differentiation, possibly through direct binding to common DNA elements.

**Figure 7.**
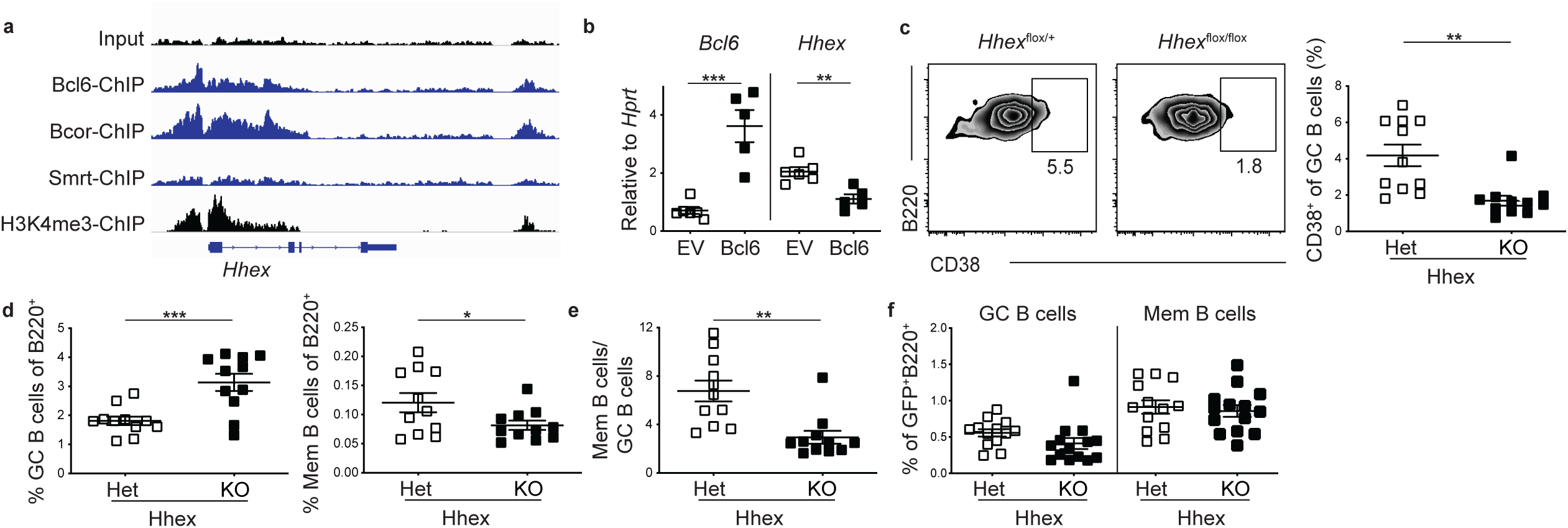
Hhex promotes the development of PreMem B cells. (**a**) Bcl6, Bcor, Smrt, and input ChIP tracks for *Hhex* from human GC B cells. H3K4me3 ChIP track from human B cells is shown to indicate promoter location. (**b**) Expression of *Bcl6* and *Hhex* relative to *Hprt* in FO B cells overexpressing *Bcl6*. Data are pooled from 2 independent experiments with 2-3 mice per group. (**c**) Representative FACS plots (left) and percentage (right) of CD38^+^ cells among splenic GC B cells (B220^+^IgD^lo^GL7^+^CD95^+^Eprhinb1^+^Tomato^+^) in *Hhex*^flox/+^ (Het) and *Hhex*^flox/flox^ (KO) S1pr2-ERT2cre TdTomato mice at day 11 post LCMV infection. (**d**) Percentage of B cells that are GC B cells (B220^+^IgD^lo^GL7^+^CD95^+^Tomato^+^) (left) and MBCs (B220^+^IgD^lo^GL7^-^ CD38^+^Tomato^+^) (right) in Hhex Het and KO mice at day 11 post LCMV infection. (**e**) Ratio of MBCs to GC B cells in Hhex Het and KO mice at day 11 post LCMV infection. Data in c-e are pooled from 3 independent experiments with 3-4 mice per group. (**f**) Percentage of GFP^+^ B cells that are MBCs (B220^+^IgD^lo^GL7^-^CD38^+^CD95^+^CD73^+^) in *Hhex*^flox/+^ and *Hhex*^flox/flox^*Ubc*^Cre – ERT2^Rosa^mTmG^ mice at day 80 post LCMV infection. Mice were treated with Tm beginning at day 35 p.i. Data are pooled from 2 independent experiments with 6-9 mice per group. Statistical analyses were performed using the unpaired two-tailed Student’s *t*-test (*, *p* < 0.05; **, *p* < 0.01; ***, *p* < 0.001). See also Figure S7.

### Hhex promotes the development of PreMem B cells

To determine whether Hhex promotes the development of PreMem B cells, we analyzed Tm treated *Hhex*^f/f^ S1pr2-ERT2creTdTomato mice at day 11 post LCMV infection. We found that Hhex KO mice had a significant reduction in PreMem B cells, identified based on previous work as the percentage of GC B cells (identified as B220^+^IgD^lo^GL7^+^CD95^+^Ephrinb1^+^Tomato^+^) expressing CD38 (**Fig. 7c, gating shown in Fig. S7a**) ^16^. We also found that while there was a decrease in the percentage of MBCs at day 11 in Hhex-deficient cells, there was also an increase in the percentage of GC B cells (**Fig. 7d, e**). Considering that there is no difference in the number of GC B cells present in Hhex KO mice at day 15 or 45 p.i. (**Fig. 6f**), the increase in GC B cells at day 11 might reflect cells that normally would transition into MBCs but rather are being transiently maintained in the GC.

To evaluate whether Hhex regulates MBC maintenance, we crossed *Hhex*^flox/flox^ and *Ubc*^Cre – ERT2^Rosa^mTmG^ mice. We then infected *Hhex*^flox/flox^*Ubc*^Cre –ERT2^Rosa^mTmG^ and control mice with LCMV. At day 35 p.i. the mice were treated with Tm to ablate Hhex expression in all cells. MBCs largely develop by day 16 post LCMV infection, so loss of Hhex after day 35 p.i. is expected to have a negligible impact on the development of MBCs (**Fig. S7b, c**). Mice were then analyzed at day 80 p.i. and the percentage of Cre expressing (GFP^+^) MBCs was determined. We found that there was no difference in the percentage of MBCs present in mice after Hhex ablation, indicating that Hhex does not regulate MBC maintenance (**Fig. 7f**).

### Hhex enhances MBC development through induction of Bcl2

To further probe how Hhex regulates MBC development we performed RNAseq analysis on Hhex-deficient MBCs (B220^+^IgD^lo^GL7^-^CD38^+^Tomato^+^ cells) at day 11 post LCMV infection. We found that there were 149 differentially expressed genes (DEGs) between Hhex KO and Het MBCs, with 82 upregulated and 67 downregulated genes in the Hhex-deficient MBCs. DEGs were defined as genes with a p_adj_<0.1 and base count >100. *Bcl6* was upregulated in Hhex-deficient cells with differential expression of Bcl6 regulated genes including *Gpr183*, *S1pr1*, *Aicda*, and *Bcl2* also being observed (**Fig. 8a**). *Bcl6* expression in Hhex-deficient MBCs was still markedly reduced from that found in GC B cells (**Fig. S8**). Hhex-deficient MBCs displayed reduced expression of the TF *Ski*, which is highly expressed in MBCs ^16^ (**Fig. 8a**). Notably, *Ski* is also reduced in Hhex-deficient lymphoid precursor cells ^36^.

**Figure 8.**
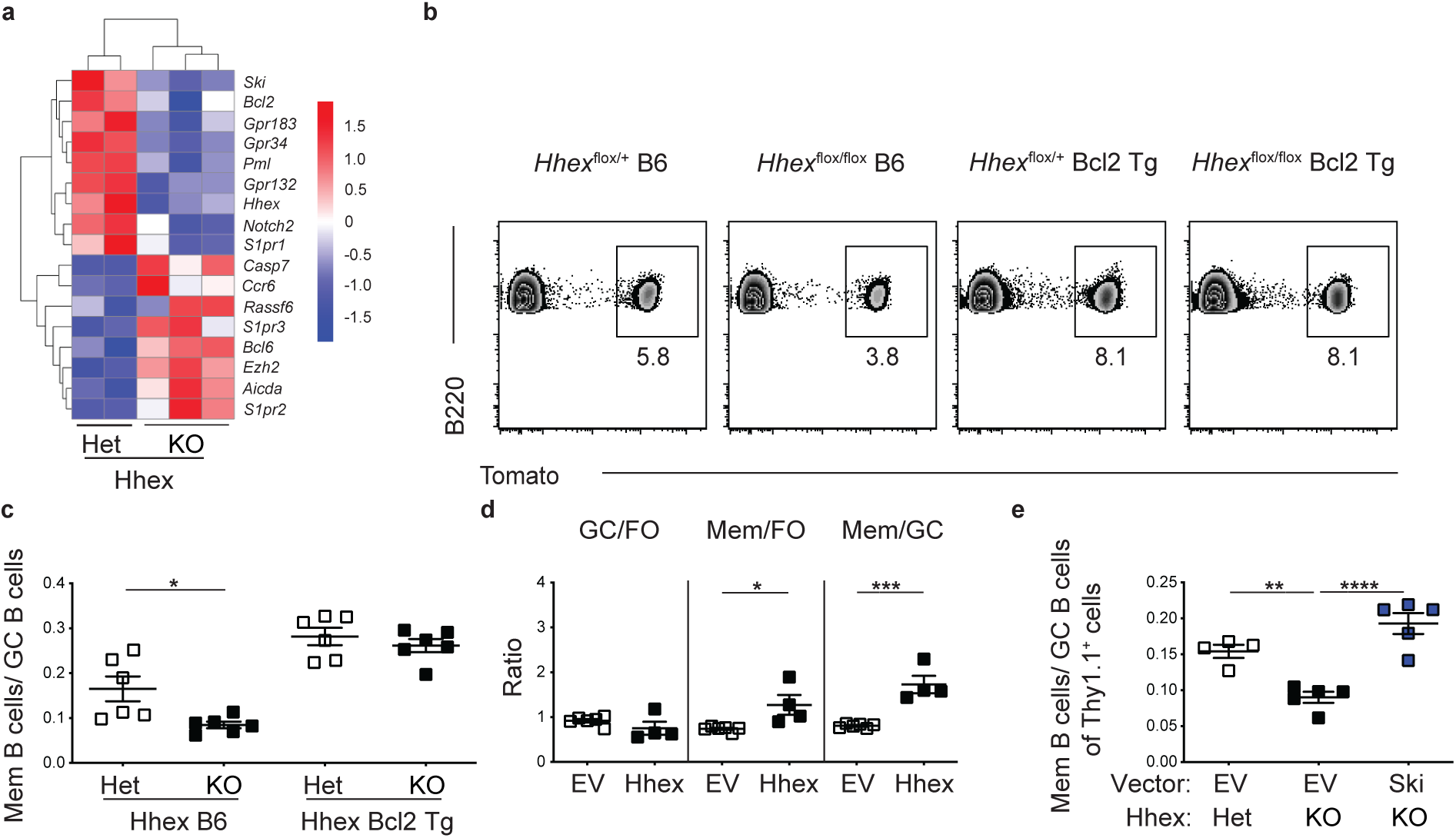
Hhex promotes MBC development through induction of *Bcl2* and *Ski*. (**a**) Heatmap of select genes from RNAseq analysis of MBCs (B220^+^IgD^lo^GL7^-^CD38^+^Tomato^+^) from *Hhex*^flox/+^ (Het) and *Hhex*^flox/flox^ (KO) S1pr2-ERT2creTdTomato mice at day 11 post LCMV infection. Data are from 3 independent experiments with 3-4 mice per experiment pooled for each sample. (**b**) Representative FACS plots of the percentage of Tomato^+^ cells among splenic B220^+^IgD^lo^GL7^-^CD38^+^ MBCs in Hhex Het and KO B6 and Bcl2-Tg S1pr2-ERT2creTdTomato mice at day 30 post LCMV infection. (**c**) Ratio of MBCs to GC B cells (B220^+^IgD^lo^GL7^+^CD95^+^Tomato^+^) in Hhex Het and KO B6 and Bcl2-Tg S1pr2-ERT2creTdTomato mice at day 30 post LCMV infection. Data in b-c are pooled from 2 independent experiments with 3 mice per group. (**d**) Ratio of transduced GC to FO B cells (left), MBCs to FO B cells (middle), and MBCs to GC B cells (right) in Hhex-overexpressing Bcl2-Tg bone marrow chimeras at day 30 post LCMV infection. Data are representative of 2 independent experiments with 4-6 mice per group. (**e**) Ratio of MBCs to GC B cells among empty vector or Ski-overexpressing Hhex Het or KO bone marrow chimeras at day 30 post LCMV infection. Data are representative of 2 independent experiments with 4-5 mice per group. Statistical analyses were performed using the unpaired two-tailed Student’s *t*-test or the ordinary one-way ANOVA with Dunnett multiple comparison testing (*, *p* < 0.05; **, *p* < 0.01; ***, *p* < 0.001; ****, *p* < 0.0001). See also Figure S8.

Bcl2 promotes memory B cell development through repression of GC B cell apoptosis ^37, 38^. To investigate whether the reduced *Bcl2* expression in Hhex-deficient MBCs contributes to impairment in differentiating into MBCs we crossed *Hhex*^f/f^ S1pr2-ERT2creTdTomato mice to Bcl2-Tg mice, a mouse line that overexpresses Bcl2 selectively in B cells ^39^. Hhex Het and KO Bcl2-Tg mice were then infected with LCMV, treated with Tm, and analyzed at day 30 p.i. We found that Hhex KO Bcl2-Tg mice displayed comparable development of MBCs relative to their Hhex Bcl2-Tg counterparts (**Fig. 8b, c**). These data suggest that Hhex promotes MBC development at least in part through induction of Bcl2, possibly via repression of Bcl6. Given that MBC maintenance is not impaired in the absence of Hhex it is likely that the Bcl2 expression in these cells, which is still markedly elevated relative to GC B cells, is sufficient for cell survival but insufficient to facilitate initial GC B cell survival and differentiation into PreMem B cells (**Fig. 7f, Fig. S8**).

### Hhex enhances MBC development through induction of the transcription factor Ski

It is likely that Hhex mediates MBC development through pathways besides Bcl2. Indeed retroviral overexpression of Hhex was still sufficient to bolster MBC development in Bcl2-Tg cells (**Fig. 8d**). Given that Hhex-deficient MBC have reduced expression of the TF Ski and that Ski overexpression was sufficient to promote MBC development (**Fig. S5a, b**), we next explored whether Hhex promotes MBC development through induction of Ski. We retrovirally overexpressed Ski in *Hhex*^f/f^ S1pr2-ERT2creTdTomato bone marrow and infected these mice with LCMV following reconstitution. Tm-treated mice were then analyzed at day 30 p.i. We found that Ski overexpression in Hhex-deficient cells was sufficient to rescue MBC development (**Fig. 8e**). These data indicate that Hhex induction of Ski expression is another mechanism by which Hhex promotes MBC development.

## DISCUSSION

Understanding of the transcriptional networks regulating MBC development is critical for efforts to design vaccines better able to induce protective humoral immunity. Here, we find that the TF Hhex cooperates with the corepressor Tle3 to promote GC B cell differentiation into MBCs. We also revealed roles for other TFs including Ski, Runx3, and Klf2 in regulating MBC differentiation. Our approach to inducibly ablate or overexpress genes specifically in GC B cells serves as a powerful tool for deciphering the signals regulating GC B cell differentiation in an *in vivo* setting.

Promotion of MBC development by Hhex is dependent on direct binding to target DNA. Tle3 is a transcriptional corepressor that also promotes MBC development but lacks DNA-binding domains. Rather, Tle3 functions through interaction with DNA-binding TFs and histone deacetylases (HDACs), which regulate histone acetylation to mediate gene repression ^33, 40^. Tle proteins can directly interact with Hhex to regulate gene repression ^18^. An Hhex mutant lacking DNA binding activity inhibits wild-type Hhex activity by sequestering Tle proteins from the nucleus ^41^. We find that Tle3 can interact with the N-terminal domain of Hhex. Further, we find that retroviral overexpression of an Hhex mutant lacking DNA binding impairs MBC differentiation, possibly via sequestration of Tle3. Tle proteins can interact with a number of TFs expressed by lymphocytes including Tcf1, Lef1, Runx1, and Runx3 ^33^. Our finding that ablation of Runx3 in GC B cells leads to enhanced MBC development supports a model in which competition for Tle3 by Hhex and other TFs, such as Runx3, serves as an important mechanism regulating MBC development.

Hhex is a homeodomain TF that is highly expressed in FO and MBCs while being downregulated in GC B cells. We find that Bcl6 directly represses the expression of Hhex, and reciprocally that overexpression of Hhex is sufficient to restrain the development of GC B cells. Moreover, newly generated Hhex-deficient MBCs have increased expression of *Bcl6* and altered expression of Bcl6 target genes. These data support a model in which Bcl6 and Hhex function in opposition, potentially by inhibiting each other, to regulate GC B cell differentiation. LZ GC B cells receiving strong positive selection (CD40 and BCR signals) will induce Myc and mTOR, and migrate to the DZ in a Foxo-1 dependent manner, where they will divide and undergo somatic hypermutation ^42–45^. LZ GC B cells that do not receive positive selection signals will remain Bcl6^hi^ and undergo apoptosis ^46^. We hypothesize that LZ GC B cells that receive low to intermediate levels of positive selection signals will downregulate Bcl6 without inducing Myc, thereby removing Bcl6-dependent repression of Hhex. As Hhex is expressed it may further suppress Bcl6, allowing for the development of PreMem B cells that are capable of migrating out of the GC and completing their differentiation into MBCs. This hypothesis is supported by work showing that GC B cells receiving low T cell help are predisposed to enter the memory pool ^14^.

Currently the extracellular signals promoting MBC development within the GC are unclear. T_FH_ cell-derived IL-9 has been suggested to promote MBC development ^12^. However, subsequent studies revealed only a modest role for IL-9 in promoting MBC development and found that 0.1% of T_FH_ cells produce IL-9 following immunization ^13^. Ingenuity Upstream Regulator Analysis found that downregulation of Bcl6 was one of the primary drivers of the differences in gene expression between GC B cells and PreMem B cells ^16^. CD40 signaling transiently downregulates Bcl6 expression in GC B cells in an NF-κB dependent manner ^47–49^. BCR signaling also promotes Bcl6 downregulation in a mitogen-activated protein kinase (MAPK) dependent manner ^50^. The T_FH_ cell cytokines IL-4 and IL-21 counteract BCR-mediated degradation of Bcl6 and help maintain high Bcl6 expression in GC B cells ^51^. Together, these studies suggest that a low level of CD40 and/or BCR-dependent signaling in GC B cells together with minimal IL-4 or IL-21, may promote MBC development through downregulation of Bcl6 and subsequent expression of Hhex.

Hhex promotes MBC development through multiple mechanisms. We find that newly formed Hhex-deficient MBCs have reduced expression of *Bcl2* and that Bcl2 overexpression in Hhex-deficient cells is sufficient to rescue MBC development. Bcl2 promotes MBC development through inhibition of apoptosis ^37, 38^. Bcl6 represses expression of Bcl2 in GC B cells by directly binding to the Bcl2 promoter and suppressing Miz1 induced activation of Bcl2 ^52^. Inadequate repression of *Bcl6* in Hhex-deficient GC B cells may therefore result in impaired Bcl2 upregulation and a reduced ability for GC B cells receiving low T cell help to survive and differentiate into MBCs. Mature Hhex-deficient MBCs still exhibit elevated Bcl2 expression relative to GC B cells and do not display impaired maintenance, indicating that Hhex is not required for the survival of MBCs once they exit the GC state.

Another gene regulated by Hhex in MBCs was the transcriptional repressor *Ski*. Hhex-deficient lymphoid progenitor cells also have reduced *Ski* expression, supporting that notion that Hhex may be a direct regulator of Ski expression ^28^. Interestingly, our findings indicate that while overexpression of Ski was sufficient to rescue MBC development in Hhex-deficient cells, ablation of Ski in GC B cells did not reduce MBC development. These data suggest that the transcriptional regulatory functions of Ski may be partially redundant with those of Hhex.

In summary, our work has provided important insight into the transcriptional circuitry governing MBC development. Moving forward it will be necessary to comprehensively define the genes directly targeted by the Hhex-Tle3 complex and to understand how TFs, such as Ski or Runx3, contribute to MBC development. Understanding of the TFs and upstream regulators governing MBC development will facilitate the design of vaccines better able to modulate these signaling pathways to promote MBC development and elicit long-lasting protective immunity.

## ACKNOWLEDGEMENTS

We thank A. Marson for the pTR plasmid, T. Okada for S1PR2-CreERT2 mice, J. An for expert technical assistance, J. Derisi for support and P.S. Jayaraman for helpful discussions. This work was supported by grants from the NIH (R01AI045073 [J.G.C], T32AI07019 [B.L.]). B.J.L. is a Howard Hughes Medical Institute Fellow of the Damon Runyon Cancer Research Foundation (DRG-2265-16). J.G.C. is an investigator of the Howard Hughes Medical Institute.

## AUTHOR CONTRIBUTIONS

Conceived and designed the experiments: B.J.L., J.G.C. Performed the experiments: B.J.L., L.D., Y.X., S.E.V. Analyzed the data: B.J.L., J.G.C. Wrote the manuscript: B.J.L., J.G.C.

## METHODS

### Mice

Adult C57BL/6 CD45.1^+^ (stock number 564) mice at least 6 weeks of age were purchased from the National Cancer Institute (NCI) or NCI at CRV. S1PR2-ERT2-Cre mice were provided by T. Okada, RIKEN Center for Integrative Medicine ^14^. *Ubc*^Cre–ERT2^ and Rosa^mTmG^ mice were provided by M. Krummel, UCSF ^53, 54^. Rosa26-LSL-Cas9 (026175), *Cg1*^Cre^ (010611), Bcl2-Tg (002321), and *Hhex*^flox^ (025396) mice were purchased from Jackson laboratories ^24, 25, 39, 55^. Mice were housed in a specific pathogen-free environment in the Laboratory Animal Research Center at the University of California, San Francisco (UCSF), and all animal procedures were approved by the UCSF Institutional Animal Care and Use Committee.

### Infections, immunizations, adoptive transfers, and treatments

Mice were infected with 2×10^5^ plaque-forming units of LCMV Armstrong administered i.p. Mice were immunized with 100µg NP-(20-29)-CGG (Biosearch technologies) mixed 1:1 with alum (Aldygrogel) for a total of 200µl volume. Tamoxifen (Sigma) was dissolved in Corn Oil (Sigma) at 20mg/ml and injected at 2mg/20g mouse i.p.. TAM diet (Envigo) containing chow replaced normal chow on the day after the final Tm dose when indicated. To overcome initial taste aversion, additional crushed TAM diet was placed in the cage at the time of first TAM diet feeding. For adoptive transfer experiments, B and T cells were enriched from LCMV immune splenocytes through EasySep (Stem Cell) negative enrichment (using CD138, NK1.1, and CD8-Biotin antibodies) and approximately 5x10^6^ cells were transferred per recipient mouse.

### Bone marrow chimeras

WT CD45.1 mice were lethally irradiated with 1,100 rads gamma-irradiation (split dose separated by 3 hours) and then i.v. injected with relevant bone marrow cells. Bone marrow was harvested by flushing the tibia and femurs of donor mice.

### Retroviral constructs and transductions

Murine Hhex, Tle3, Runx3, Ski, Mndal, Zmynd11, Klf2, and Klf3 retroviral constructs were made by inserting the mouse open reading frame into the MSCV2.2 retroviral vector followed by an internal ribosome entry site (IRES) and Thy1.1 as an expression marker. *Hhex* point mutations were introduced by PCR followed by assembly using NEBuilder HiFi DNA Assembly Master Mix (New England BioLabs). *Hhex* truncation was introduced by PCR. Loxp-EGFP-loxp was also cloned into the MSCV2.2 retroviral vector followed by an internal ribosome entry site (IRES) and Thy1.1 as an expression marker. The murine *Hhex* open reading frame was then inserted the Loxp-EGFP-IRES-Thy1.1 MSCV2.2 vector. For the TF screen, sgRNAs were cloned into the pTR-MSCV-IRES-Thy1.1 or pTR-MSCV-IRES-BFP vector. sgRNA sequences were selected using the Broad Institute sgRNA Designer cross referenced with the Benchling Crispr Guide tool. For sgRNAs that did not begin with a G, a G was added. A primer with the sequence GTGGAAAGGACGAAACACC-sgRNA sequence-GTTTTAGAGCTAGAAATAG was then order and cloned into the pTR-MSCV-IRES vector using the NEBuilder HiFi DNA Assembly Reaction protocol (New England BioLabs). Transduction efficiencies for all experiments varied between constructs and sets of bone marrow chimeras.

The following sgRNA were used:

*Hhex* sgRNA 1: GATACAGCGGGACTCCCACGG; sgRNA 2: GTCGACGACATCTTGGGTCGC *Foxp1* sgRNA 1: GTGGTCTCTCGGTGCAAACTA; sgRNA 2: GCAGCTGCACGCTGTGCTCG *Ski* sgRNA 1: GATGGTGCGATCGGGCTCCCC; sgRNA 2: GCTCCCGCTCCCGTGCGCCC *Klf2* sgRNA 1: GTGAGGACCTAAACAACGTGT; sgRNA 2: GTCCATGGGATTGGACGGTCT *Klf3* sgRNA 1: GACCCCACCCGCGCGTCCGTG; sgRNA 2: GTCTGGGCAGTCACATGACGC *Tle3* sgRNA 1: GCCGGGATTTAAATTCACTG; sgRNA 2: GCTTGCGGATACATGGCAGGG *Foxo3* sgRNA 1: GCCCGAGAGTCCCCTCGTCG; sgRNA 2: GCAGCATGGCCGAATCCTCG *Bhlhe41* sgRNA 1: GCGCAGTGCGTGTGTGCGCG; sgRNA 2: GAGGGCGAGCGAGCGAGCACG *Runx3* sgRNA 1: GCAGAGTATCATTAAATGGT; sgRNA 2: GTTTGTGGCTAGACATTCCTG *Zeb2* sgRNA 1: GTAACACGTCAGTCCGTCCCC; sgRNA 2: GACGCGCCACCTATCTTTGTG *Jun* sgRNA 1: GACGGTCCGTCACTTCACGCG; sgRNA 2: GACTCCGCTAGCACTCACGT *Pml* sgRNA 1: GCAGAGTCCTGTTGTCGACA; sgRNA 2: GCTCGAAGAACACGTTATCCA *Bhlhe40* sgRNA 1: GTGGCATCCCAGCGCATTGCA; sgRNA 2: GAGGATCCGAGGGTCTCAAG *Tcf4* sgRNA 1: GATTATCAATGTGACTCCTCG; sgRNA 2: GATTGTAATCCATTCACATCC *Zfp182* sgRNA 1: GCTACATCCTCAAATGTCAC; sgRNA 2: GTGTAGCTGTGGATTTCACTC *Mndal* sgRNA 1: GAAAGCGATCGCAAAAACTGA; sgRNA 2: GTGTCTAGTCCAGCATATTTG *Zmynd11* sgRNA 1: GATTGTCGAAACTCTAACAGT; sgRNA 2: GAATGATACACACGAAAACAC *Sub1* sgRNA 1: GTGGTGAGACTTCTAGAGCAC; sgRNA 2: GTCAGTGTTCGGGACTTCAA *Arid5a* sgRNA 1: GATAGTGGCGGCGTGTGCAT; sgRNA 2: GAGCGACACACGCCCATCGAG *Cd38* sgRNA 1: GCGCCTTGGTAGTAGGGATCG; sgRNA 2: GAGCCCAGATCGGTCTCGGAG *Cxcr3* sgRNA 1: GTGAGGGCTACACGTACCCGG; sgRNA 2: GTAGCACCACCAGGTGATAGG Control sgRNA: GCGAGGTATTCGGCTCCGCG

Retrovirus was generated by transfecting PLAT-E packaging cell line with 10 µg plasmid DNA and 10 mg Lipofectamine 2000 (Fischer). For transduction of bone marrow, WT, Bcl2-Tg, *Hhex*^flox/+^ S1pr2-ERT2creTdTomato, *Hhex^f^*^lox/flox^ S1pr2-ERT2creTdTomato, Rosa26-LSL-Cas9^f/+^ *Cg1*^Cre/+^ were injected i.v. with 3 mg 5-fluorouracil (Sigma). Bone marrow was collected after 4 days and cultured in DMEM containing 15% FBS, antibiotics (penicillin (50 IU/mL) and streptomycin (50 mg/mL); Cellgro) and 10 mM HEPES, pH 7.2 (Cellgro), supplemented with IL-3, IL-6 and stem cell factor (at concentrations of 20, 50 and 100 ng/mL, respectively; Peprotech). Cells were ‘spin-infected’ twice at days 1 and 2 and then transferred into irradiated recipients.

### Antibodies for flow cytometry staining

Spleens were mashed through a 70µm cell strainer, and red blood cells were lysed with RBC lysing buffer. Lymphocytes were then washed and counted. The following antibodies were used for flow cytometry and microscopy staining: Phycoerythrin (PE) anti-CD86 (105008), phycoerythrin–indotricarbocyanine (PE-Cy7) anti-CD38 (102718), A647 anti-CD38 (102716), Brilliant Violet 605 (BV605) anti-CD45.1 (110738), allophycocyanin (APC) anti-GL7 (144606), Pacific Blue (PacBlue) anti-GL7 (144614), peridinin chlorophyll protein Cy5.5 (PerCpCy5.5) anti-CD73 (127214), PacBlue anti-IgD (405712), PerCpCy5.5 anti-IgD (405710), APC anti-CD80 (104718), PE anti-CD11b (101208), PE anti-CD44 (103008), PE anti-IgD (405705), APC Cy7 anti-B220 (103224), FITC anti-GL7 (144604), PE anti-CD138 (142504), BV412 anti-CD138 (142506), BV711 anti-Thy1.1 (202539), Alexa647 anti-Thy1.1 (202508), PerCpcy5.5 anti-Thy1.1 (202516), A647 anti-Bcl-2 (633510) (all from BioLegend); APC anti-CXCR4 (558644), PE anti-CD95 (554258), PE Cy7 anti-CD95 (557653), APC anti-Bcl6 (561525), APC anti-CCR6 (557976), Biotin anti-CD35 (553816), Biotin anti-CXCR4 (551968), APC anti-TCRb (17-5961-82), BV605 Streptavidin (563260), APC Cy7 anti-CD19 (115530), FITC Rabbit anti-active Caspase3 (560901), A647 anti-Bcl-6 (561525) (all from BD Biosciences); PerCPCy5.5 anti-CD45.2 (65-0454-U100) (Tonbo Biosciences); Biotin goat anti-mouse polyclonal Ephrin-B1 (BAF473) (R&D Systems); FITC anti-BrDU (11-5071-42) ; Fixable viability dye eFluor780 (65-0865-18) (eBiosciences). Flow cytometry data were acquired on a BD LSRII with FACSDiva software and were analyzed with FlowJo software (TreeStar).

### RNAseq library preparation and data analysis

Total RNA was purified from FACS sorted cells using the RNeasy Micro kit (Qiagen). RNA quality was assessed with an Agilent 2100 Bioanalyzer (RNA integrity number>9 for all samples). Barcoded sequencing libraries were generated with 100ng of RNA with an Ovation RNAseq System V2 (Nugen), KAPA Hyper Prep Kit for Illumina (KAPA Biosystems), and NEXTflex DNA barcodes (Bioo Scientific). Single-end sequencing was performed on an Illumina HiSeq 4000 (UCSF Center for Advanced Technology) and sequences reported as FASTQ files, which were aligned to the mm10 genome with STAR (Spliced Transcript Alignment to a Reference). Mappable reads were counted with HTseq and imported into RStudio software for analysis of differential expression with DESeq2 software.

### Droplet based single-cell RNA sequencing

Immediately post-sorting, the indicated B cell populations were run on the 10X Chromium (10X Genomics) and then through library preparation by the Institute for Human Genetics at UCSF following the recommended protocol for the Chromium Single Cell 3^’^ Reagent Kit (v3 Chemistry) ^56^. Libraries were run on the HiSeq4000 for Illumina sequencing. Post-processing and quality control were performed by the Genomics Core Facility at the Institute for Human Genetics at UCSF using the 10X Cell Ranger package (v3.0.2, 10X Genomics). Reads were aligned to mm10 reference assembly and analyzed using the Seurat and Monocle software packages.

### Expression and purification of GST fusion proteins

Mouse GST-Hhex fusion proteins (containing the N-terminal domain) were generated using the GST Gene Fusion System (GE), according to manufacturer’s instructions. Briefly, gene segments encoding amino acids 1-141 of Hhex PCR amplified from a wild type or F33E mutant template were cloned into the pGEX-2T vector and then expressed in BL21 pLysS cells. Fusion protein expression was induced with 1 mM isopropyl-1-thio-β-D-galactopyranoside. Cells were harvested and lysed by sonication in PBS containing 1% Triton X-100, 1mM EDTA, and protease inhibitors (aprotinin, leupeptin, PMSF) GST-Hhex fusion proteins were purified over glutathione-Sepharose 4B beads (Sigma) and eluted with 10 mM glutathione according to manufacturer’s instructions. GST-Hhex fusion proteins were then assayed for purity by SDS-PAGE followed by staining with Coomassie Blue. GST has a MW of 26.9 kDa and the expected size of the fusion protein was 15.5kDa.

### ***In vitro* Binding Assays**

Tle3 containing a flag tag in the C-terminus was cloned into the pT_N_T plasmid and transcription and translation was subsequently performed using the T_N_T Quick Coupled Transcription/Translation System (Promega) according to the manufacturer’s protocol. Approximately 20µg of each GST fusion protein or an equimolar amount of GST protein as judged by Coomassie staining was incubated with 10µl of the *in vitro* translated Tle3 for 1 hour at room temperature. 25µl of glutathione-Sepharose 4B beads (Sigma), prepared according to the manufacturer’s protocol, was then added. The beads were then washed four times with binding buffer (TBS, 50mM Tris-HCl pH 7.4, 150mM NaCl). Bound proteins were eluted by boiling the beads in 2x Laemmli Sample Buffer (BIO-RAD). Samples were then loaded onto a NuPAGE 4-12% Bis-Tris Protein Gel (Thermo) and then transferred onto a nitrocellulose membrane. Membranes were stained using rabbit anti-Flag primary antibody (Cell Signaling Technologies, #14793S) followed by a donkey anti-rabbit IRDye 680 RD (Licor, #926-68073) and visualized using a Licor Odyssey CLx.

For the radioactive ligand binding assay, EasyTag L-[^35^S]-Methionine (Perkin Elmer) was added to the T_N_T Quick Coupled Transcription/Translation reaction. Buffer exchange was then performed using a NAP-5 column (GE Healthcare) and protein amount quantified using a MicroBeta TriLux liquid scintillation counter. Approximately 12µg of each GST fusion protein or an equimolar amount of GST protein as judged by Coomassie staining was incubated with the *in vitro* translated Tle3 for 1 hour at room temperature in a 96-well plate. All samples were then added to a preblocked 96-well filter plate (Corning) containing 5µl of glutathione-Sepharose 4B beads (Sigma), prepared according to the manufacturer’s protocol. Samples were incubated for 20 minutes at room temperature and then washed four times. Samples were then quantified using a MicroBeta TriLux liquid scintillation counter with counts normalized based on the amount of input protein.

### Real-time PCR

Total RNA was isolated and converted to cDNA. A StepOnePlus realtime PCR system (Applied Biosystems) with Power SYBR Green PCR Master Mix (Thermo Fisher) and the appropriate primer pairs (Integrated DNA Technologies) were used for real-time PCR. The following primers were used:

*Bcl6* Forward: TTGTCATCGTGGTGAGCCG ; Reverse: TTGCATTTCAACTGGTCAGTG *Hhex* Forward: AAATACCTCTCCCCACCCGA; Reverse: TGTTGCTTTGAGGATTCTCCTGT

### Statistical analysis

Results represent the mean ± SEM unless indicated otherwise. Statistical significance was determined by the unpaired Student’s t test or the ordinary one-way ANOVA with Dunnett multiple comparison testing. Statistical analyses were performed using Prism GraphPad software v8.0. (*, *p* < 0.05; **, *p* < 0.01; ***, *p* < 0.001; ****, *p* < 0.0001).

## DATA AND CODE AVAILABILITY

Raw and processed data files for the scRNAseq and RNAseq analysis have been deposited in the NCBI Gene Expression Omnibus under accession number GEO: pending.

## Abbreviations

GC: germinal center
MBC: memory B cell
T_FH_: T follicular helper cell
DEGs: differentially expressed genes
TF: transcription factor
scRNAseq: single cell RNA sequencing
FO: follicular
DZ: dark zone
LZ: light zone
FO: follicular
Sg: single guide
ChIP: chromatin immunoprecipitation
GST: glutathione S-transferase
HDAC: histone deacetylase
LCMV: lymphocytic choriomeningitis virus
MAPK: mitogen activated protein kinase

---

**Supplementary Figure 1.**
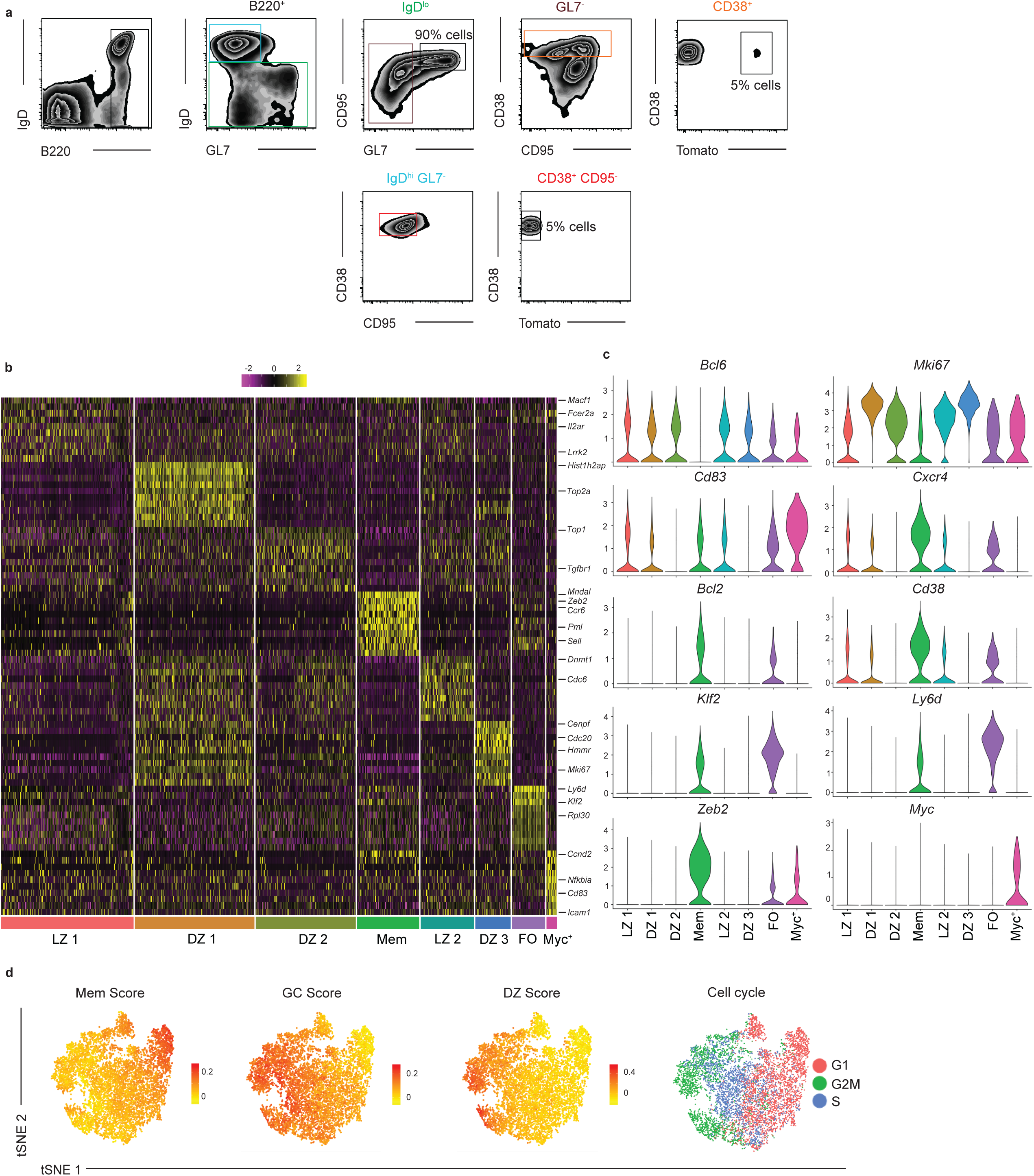
Differential gene expression for eight B cell clusters. (**a**) Gating strategy for the sorted B cells used for droplet-based scRNAseq analysis day 11 post LCMV infection. The sorted population consisted of a mixture of 90% GC B cells, 5% FO B cells, and 5% MBCs (**b**) Heatmap of each cell’s (column) expression of the top ten DEGs per cluster (rows). Select genes are labeled. Log-normalized expression scaled for each gene. Cluster name displayed below. (**c**) Violin plots of select gene expression by cluster with highest log-normalized expression value labeled. (**d**) Enrichment score for gene signature distinguishing MBCs (far left), GC B cells (middle left), and DZ GC B cells (middle right) projected onto tSNE plots. Color scaled for each gene with highest log-normalized expression level noted. Stage of cell cycle based on gene expression projected onto tSNE plot (far right).

**Supplementary Figure 2.**
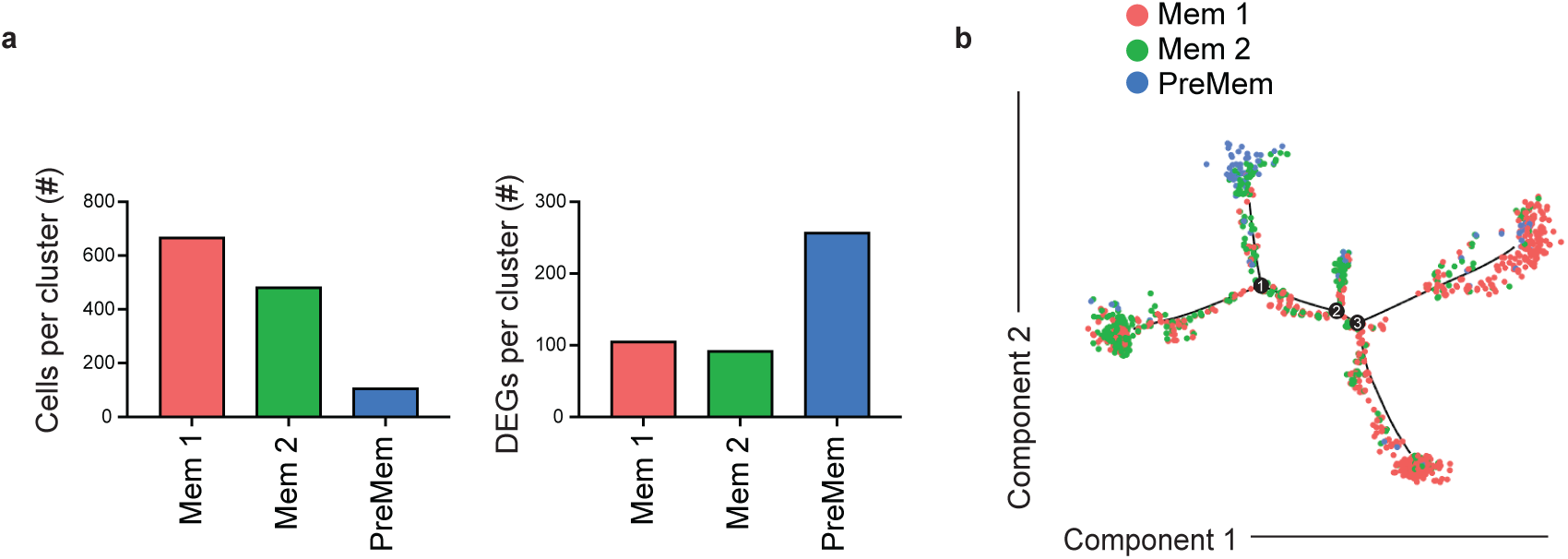
Pseudotime trajectory analysis of memory cluster cells. (**a**) Number of cells per cluster (left) and DEGs per cluster (right) following subclustering analysis of Mem cluster. (**b**) Pseudotime trajectory analysis of Mem cluster cells. Each point is a single cell colored by cluster assessment.

**Supplementary Figure 3.**
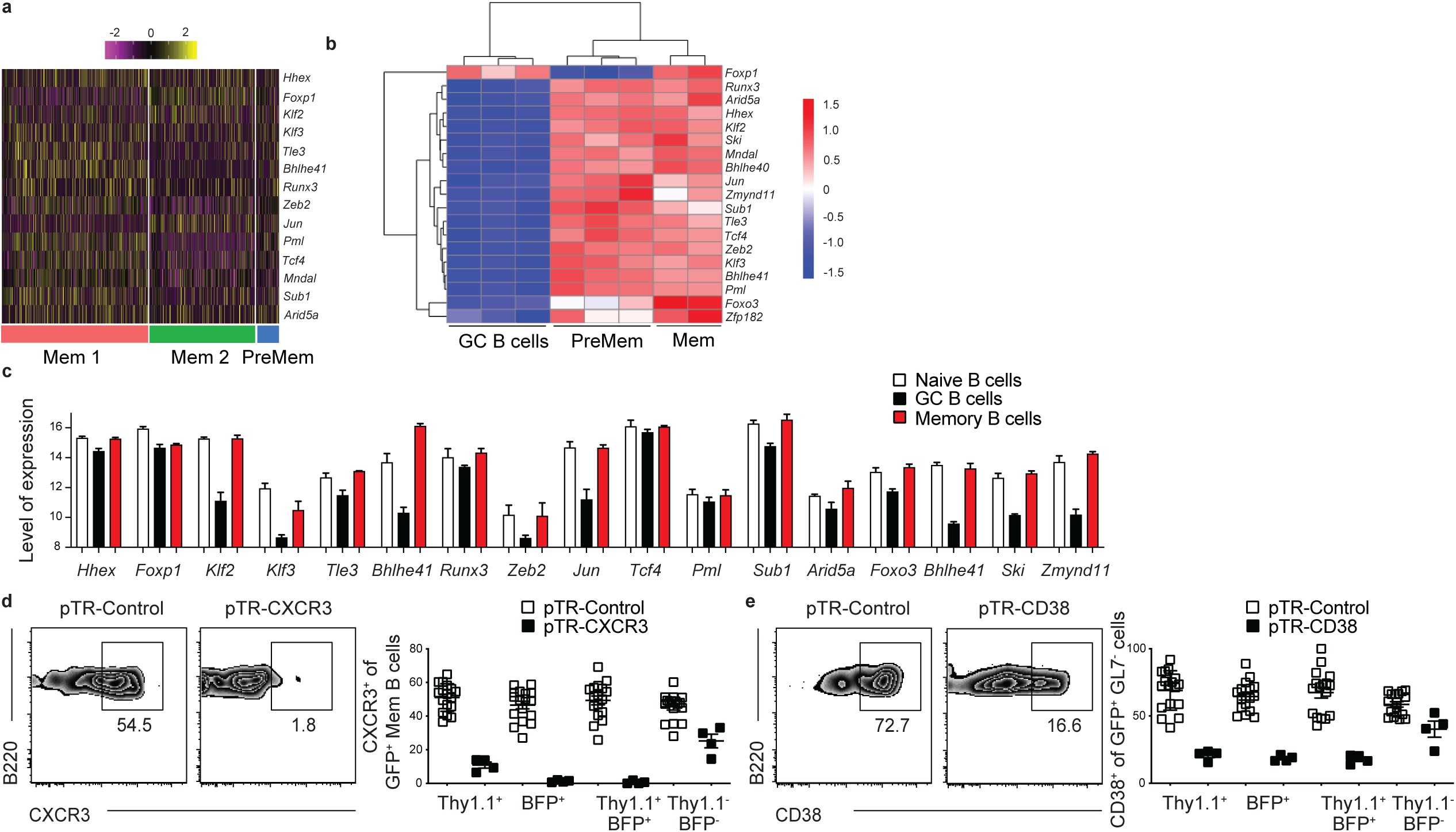
Transcription factor expression in mouse and human memory B cells. (**a**) Heatmap of each cell’s (column) expression of select TFs per cluster (rows). Log-normalized expression scaled for each gene. Cluster name displayed below. (**b**) Heatmap of TF expression from bulk RNAseq of splenic GC B cells, PreMem B cells, and MBCs at day 11 post LCMV infection. (**c**) TF expression in human naïve, GC, and MBCs as determined using datasets compiled by Genevestigator. 3 samples were included in each dataset. Level of expression is a unitless value on a log2 scale computed following normalization and scaling of raw data in order to allow comparison of expression of given gene between samples. (**d**) Representative FACS plots of CXCR3^+^ cells (left) among Thy1.1^+^BFP^+^ MBCs (B220^+^IgD^lo^GL7^-^ CD38^+^GFP^+^) at day 30 post LCMV infection in mice containing cells transduced with sgRNAs targeting a control nonexpressed gene and CXCR3. Percentage of CXCR3^+^ cells among Thy1.1^+^, BFP^+^, Thy1.1^+^BFP^+^, and Thy1.1^-^BFP^-^ MBCs (right). (**e**) Representative FACS plots of CD38^+^ cells (left) among Thy1.1^+^BFP^+^ B220^+^IgD^lo^GFP^+^ cells at day 30 post LCMV infection in mice containing cells transduced with sgRNAs targeting a control nonexpressed gene and CD38. Percentage of CD38^+^ cells among Thy1.1^+^, BFP^+^, Thy1.1^+^BFP^+^, and Thy1.1^-^BFP^-^ B220^+^IgD^lo^GFP^+^ (right).

**Supplementary Figure 4.**
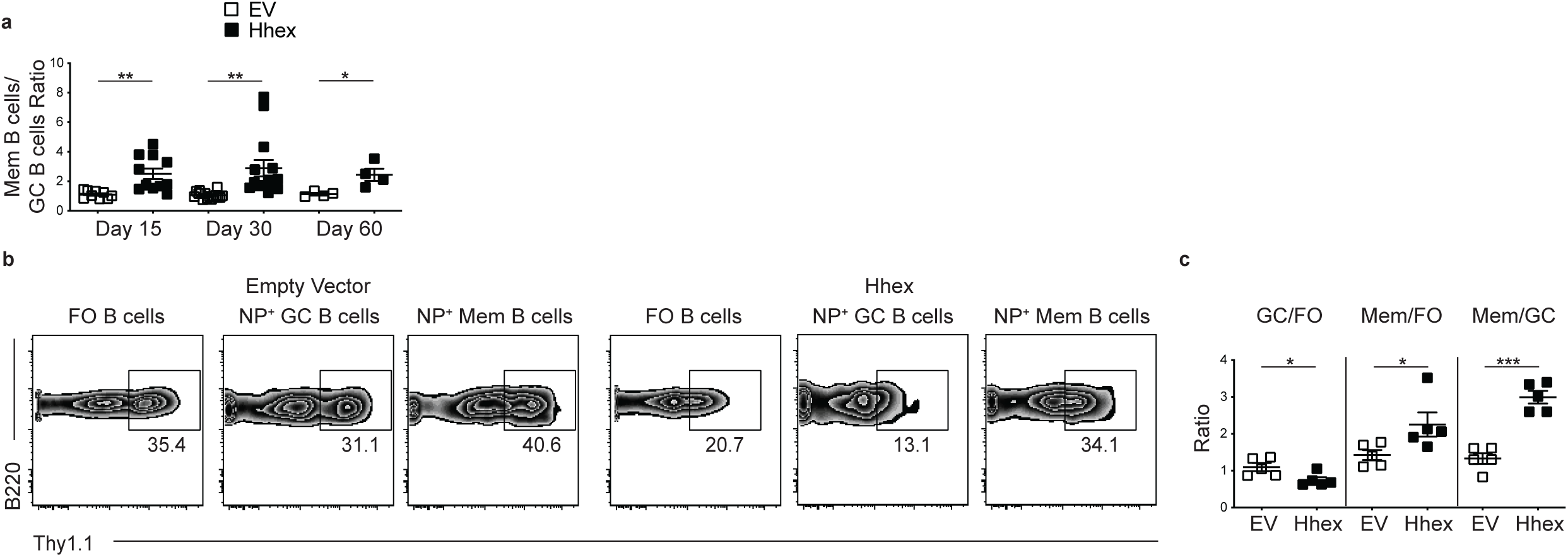
Overexpression of Hhex promotes memory B cell differentiation following Th1-type and Th2-type immunization. (**a**) Ratio of transduced (Thy1.1^+^) splenic MBCs (B220^+^IgD^lo^GL7^-^CD38^+^CD95^+^CD73^+^) to GC B cells (B220^+^IgD^lo^GL7^+^CD95^+^) in Hhex-overexpressing bone marrow chimeras at day 15, 30, and 60 post LCMV infection. Data are pooled from 8 independent experiments with 4-6 mice per group. (**b**) Representative FACS plots of the percentage of transduced (Thy1.1^+^) cells among splenic FO (B220^+^IgD^hi^GL7^-^CD38^+^CD95^-^), GC (B220^+^IgD^lo^GL7^+^CD95^+^NP^+^), and MBCs (B220^+^IgD^lo^GL7^-^CD38^+^NP^+^) at day 30 post NP-CGG in alum immunization in empty vector and Hhex-overexpressing bone marrow chimeras. (**c**) Ratio of transduced GC to FO B cells (left), MBCs to FO B cells (middle), and MBCs to GC B cells (right). Data are representative of 2 independent experiments with 4-6 mice per group. Statistical analyses were performed using the unpaired two-tailed Student’s *t*-test (*, *p* < 0.05; ***, *p* < 0.001).

**Supplementary Figure 5.**
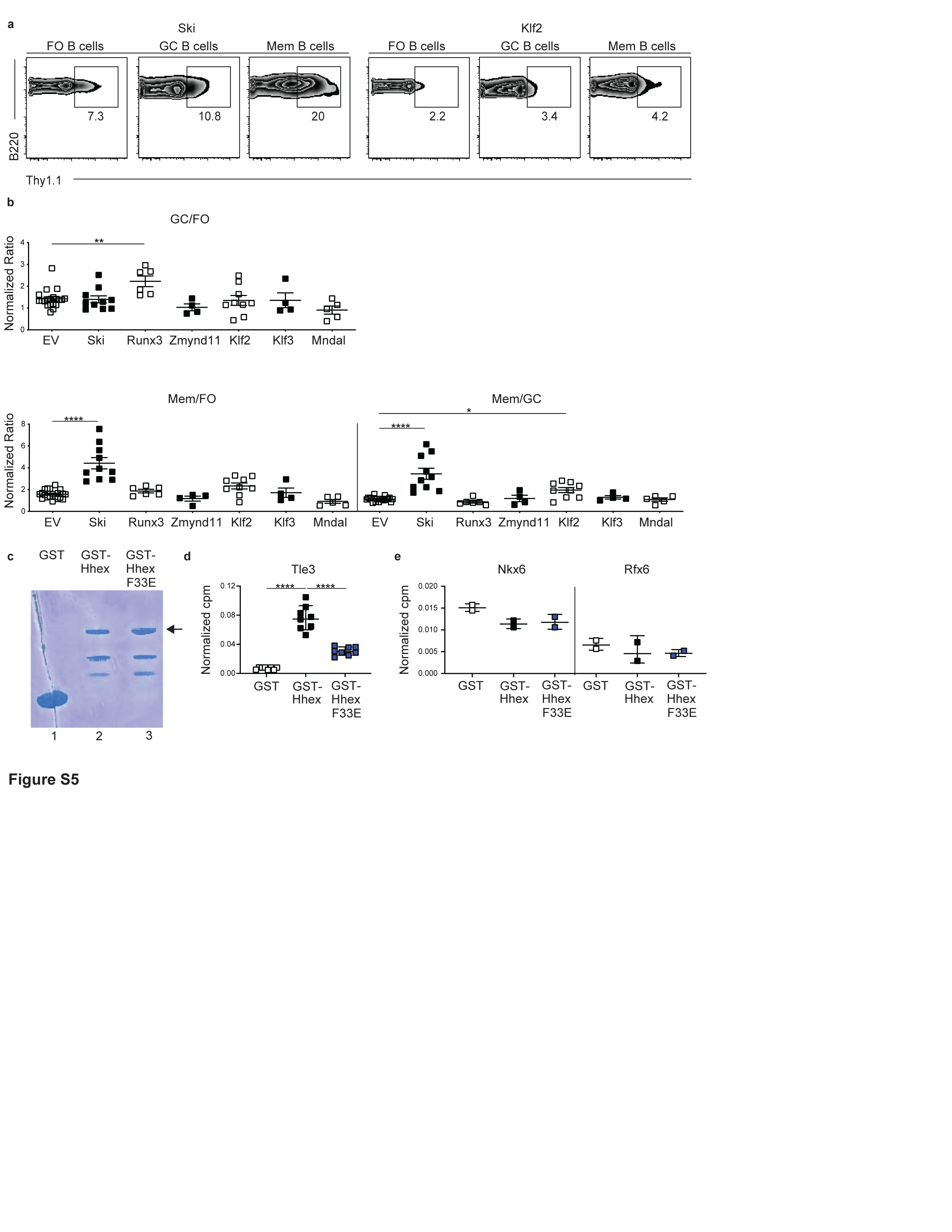
Retroviral overexpression of Ski and Klf2 promotes memory B cell differentiation. (**a**) Representative FACS plots of the percentage of transduced (Thy1.1^+^) cells among splenic FO (B220^+^IgD^hi^GL7^-^CD38^+^CD95^-^), GC (B220^+^IgD^lo^GL7^+^CD95^+^), and MBCs (B220^+^IgD^lo^GL7^-^CD38^+^CD95^+^CD73^+^) at day 30 post LCMV infection in Ski and Klf2-overexpressing bone marrow chimeras. (**b**) Ratio of transduced GC to FO B cells (top), MBCs to FO B cells (bottom left), and MBCs to GC B cells (bottom right). Vector used to transduce cells listed below. Data are pooled from 4 independent experiments with 3-6 mice per group. (**c**) Coomassie staining of gel loaded with GST (lane 1), GST-Hhex (lane 2), and GST-Hhex F33E (lane 3) fusion proteins. Arrow indicates full-length fusion protein. (**d**) Interaction of in vitro translated and transcribed Tle3 with glutathione beads coated with GST, GST-Hhex, or GST-Hhex F33E as quantified using a radioactive ligand binding assay. Counts per minute (cpm) are normalized based on the count of the input Tle3 used in the binding assay. (**e**) Interaction of in vitro translated and transcribed Nkx6 (left) and Rfx6 (right) with glutathione beads coated with GST, GST-Hhex, or GST-Hhex F33E as quantified using a radioactive ligand binding assay. Counts per minute (cpm) are normalized based on the count of the input Nkx6 or Rfx6 used in the binding assay. Statistical analyses were performed using the unpaired two-tailed Student’s *t*-test or the ordinary one-way ANOVA with Dunnett multiple comparison testing (*, *p* < 0.05; **, *p* < 0.01; ****, *p* < 0.0001).

**Supplementary Figure 6.**
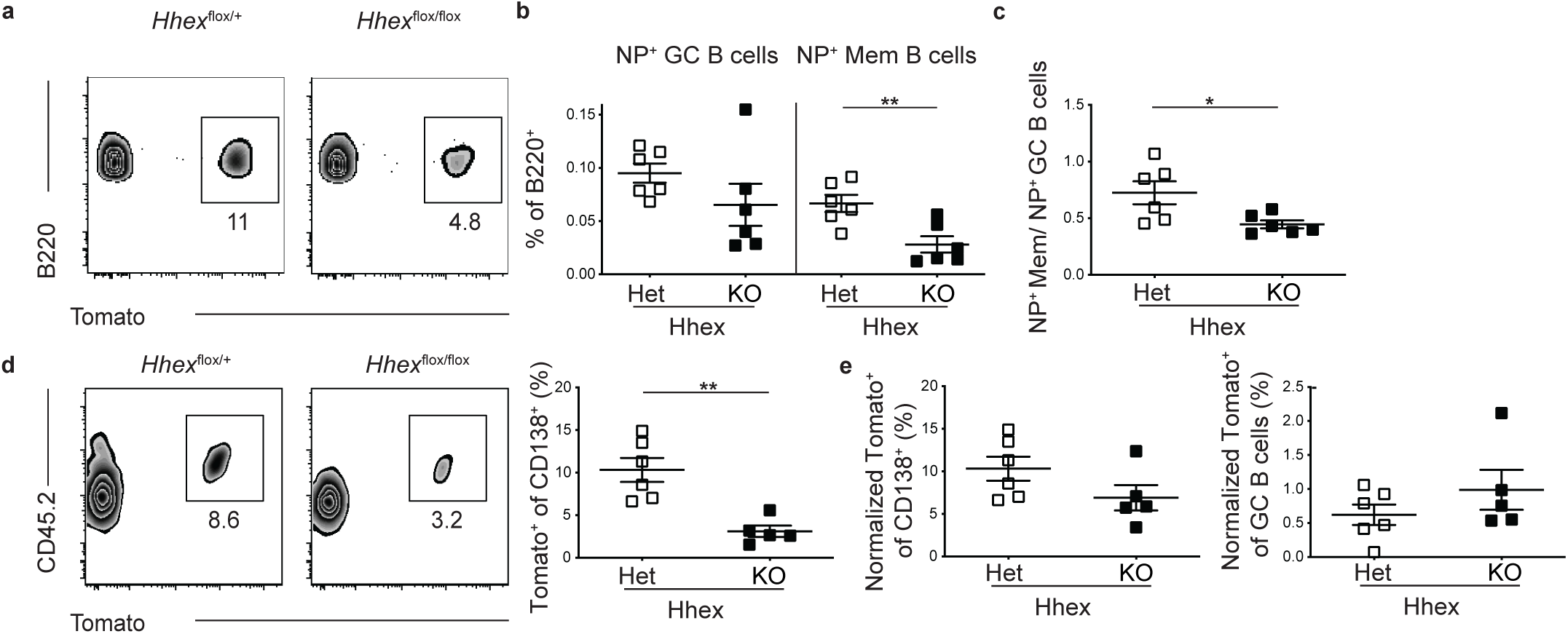
Ablation of Hhex in GC B cells impairs memory B cell differentiation following Th2-type immunization. (**a**) Representative FACS plots of the percentage of Tomato^+^ cells among splenic B220^+^IgD^lo^GL7^-^CD38^+^NP^+^ cells in Hhex Het and KO mice at day 30 post NP-CGG in alum immunization. (**b**) Percentage of B cells that are GC B cells (B220^+^IgD^lo^GL7^+^CD95^+^NP^+^Tomato^+^) and MBCs (B220^+^IgD^lo^GL7^-^CD38^+^NP^+^Tomato^+^) in Hhex Het and KO mice at day 30 post NP-CGG in alum immunization. (**c**) Ratio of NP^+^ MBCs to NP^+^ GC B cells in Hhex Het and KO mice at day 30 post NP-CGG in alum immunization. Data in a-c are pooled from 2 independent experiments with 3 mice per group. (**d**) Representative FACS plot (left) and percentage (right) of CD138^+^ cells that are CD45.2^+^Tomato^+^ at day 5 post challenge with NP-CGG in alum. Equivalent numbers of splenic CD45.2^+^ T and B cells from Hhex Het and KO NP-CGG in alum immune mice were transferred to naïve CD45.1^+^ recipients one day prior to challenge. (**e**) Percentage of CD138^+^ cells (left) and GC B cells (right) that were CD45.2^+^Tomato^+^ at day 5 post challenge with NP-CGG in alum when normalized to the percentage of MBCs present in the transferred cells. Data are pooled from 2 independent experiments with 3 mice per group. Statistical analyses were performed using the unpaired two-tailed Student’s *t*-test (*, *p* < 0.05; **, *p* < 0.01).

**Supplementary Figure 7.**
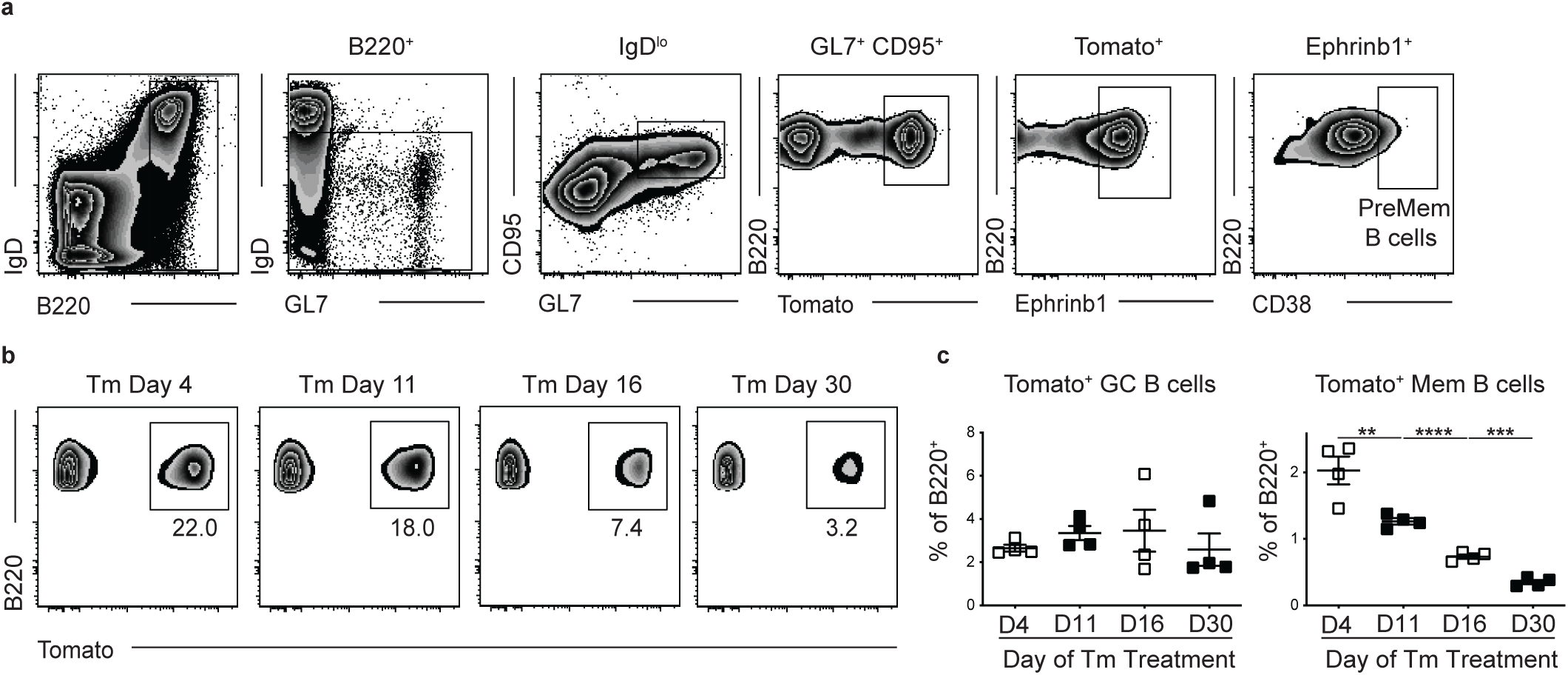
Memory B cells develop early following LCMV infection. (**a**) Gating scheme for PreMem B cells at day 11 post LCMV infection. (**b**) Representative FACS plots of Tomato expression in B220^+^IgD^lo^GL7^-^CD38^+^ cells at day 60 post LCMV infection in S1pr2-ERT2creTdTomato mice treated with Tm beginning at day 4, 11, 16, or 30 p.i. (**c**) Percentage of Tomato^+^ GC (left) and MBCs (right) at day 60 post LCMV infection in S1pr2-ERT2creTdTomato mice treated with Tm beginning at day 4, 11, 16, or 30 p.i. Data are representative of 3 independent experiments with 4-6 mice per group. Statistical analyses were performed using the ordinary one-way ANOVA with Dunnett multiple comparison testing (**, *p* < 0.01; ***, *p* < 0.001; ****, *p* < 0.0001).

**Supplementary Figure 8.**
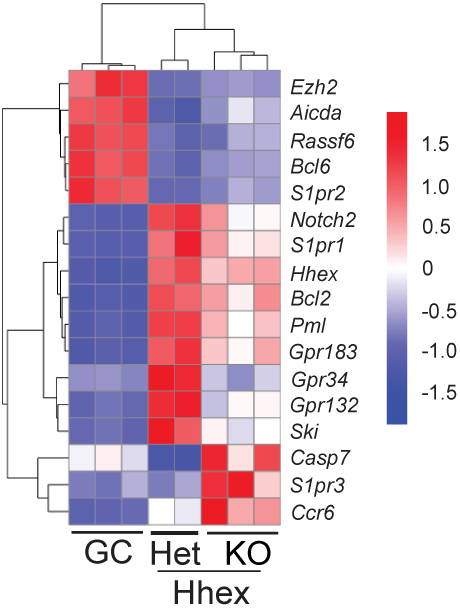
Expression of differentially expressed genes in MBCs. Heatmap of select genes from RNAseq analysis of MBCs (B220^+^IgD^lo^GL7^-^CD38^+^Tomato^+^) from *Hhex*^flox/+^ (Het) and *Hhex*^flox/flox^ (KO) S1pr2-ERT2creTdTomato mice and GC B cells (B220^+^IgD^lo^GL7^+^CD95^+^Ephrinb1^+^*S1pr2*^Venus/+^) ^16^ at day 11 post LCMV infection. Data are from 3 independent experiments with 3-4 mice per experiment pooled for each sample.

